# Master control genes in the regeneration of rod photoreceptors from endogenous progenitor cells in zebrafish retina

**DOI:** 10.1101/2025.02.03.636263

**Authors:** Eyad Shihabeddin, Abirami Santhanam, Stephan Tetenborg, Alexandra L. Aronowitz, Haichao Wei, Guoting Qin, Chengzhi Cai, Jiaqian Wu, John O’Brien

## Abstract

Retinitis Pigmentosa is a chronic retinal degenerative disease characterized by the gradual loss of rod, and later, cone photoreceptors until the individual is completely blind. Regeneration of photoreceptors from endogenous progenitor cells is a possible therapeutic approach, but mammals do not do this naturally. Mammalian models can be induced to generate retinal progenitors from Müller glial cells, but there has been limited success in rod photoreceptor specific regeneration. Unlike mammals, zebrafish have the natural ability to regenerate neurons after injury or disease and can provide insight into the molecular mechanisms of regeneration. In this study, we used a zebrafish model of Retinitis Pigmentosa to investigate the class of progenitors responsible for rod photoreceptor regeneration in the context of chronic disease. Using bioinformatic analyses of single-cell RNA sequencing datasets, we identified master regulator genes responsible for proliferation of retinal progenitors, differentiation of progenitors into rod photoreceptors, and maturation of the new rod photoreceptors. Using transient knockdown of gene expression in adult regenerating retina we determined that *e2f1*, *e2f2*, *e2f3* and *aurkb* are critical for proliferation of progenitors, and *prdm1a* is critical for differentiation of progenitors into rod photoreceptors. This study provides a list of master regulators responsible for the specific regeneration of rod photoreceptors during chronic retinal degeneration.

**Impact Statement:** Identification of master regulating genes that drive the proliferation of progenitor cells and their differentiation specifically into rod photoreceptors provides insight that can be used to develop regenerative therapies for retinal degenerative diseases.

## Introduction

Retinitis Pigmentosa (RP) is an inherited chronic retinal degenerative disease that affects about 1 in 4,000 people worldwide and results in nearly a billion dollars of added healthcare costs annually to patients in the United States (US) alone (1–3). With loss of vision starting in young adults, some patients are completely blind by their late twenties/early thirties (1). Characterized by the progressive loss of first rod photoreceptors and, later, all photoreceptors, much research has gone into investigating how to reduce progression of the disease as well as how to restore vision. While gene therapy for one variant of RP has been effective, the large number of genes that can cause RP has left many patients with no effective treatment options (4–6). As such, regenerative medicine offers significant hope for preserving and restoring vision to patients with RP (7, 8).

While mammalian models are excellent for studying the progression of retinal degeneration, they have only limited capacity to naturally regenerate neurons through Müller glial cell (MGC) activity (9–11). Zebrafish (Zf), on the other hand, have a retinal circuitry very comparable to mammals as well as the innate ability to regenerate neurons following retinal injury or disease (12, 13). In acute damage models, MGCs activate, de-differentiate into multipotent progenitor cells that then proliferate, re-differentiate into retinal cells, and integrate into the retina (10, 14–18). In chronic retinal degeneration models, such as Zf with RP, how rod photoreceptor regeneration occurs still remains largely unexplored (19). Currently, it is known that regenerated rod photoreceptors come from progenitor cells that reside in the outer nuclear layer (ONL) rather than directly from MGCs (20–23). MGCs are predominantly inactive in these models, while the progenitors in the outer nuclear layer proliferate and differentiate into rod photoreceptors (19, 20, 22–25). However, the transcription factors that regulate proliferation and differentiation of these progenitor cells remain unknown.

Previously, we have generated and characterized a P23H mutant rhodopsin transgenic zebrafish model of RP that shows continuous degeneration of rod photoreceptors and regeneration through progenitor cells found predominantly in the ONL (26). We have also previously utilized Single-Cell RNA Sequencing (SC-RNA seq) to identify the molecular basis of retinal remodeling that occurs in our Zf model (27). In this study, we identify the master regulators critical for each step of rod photoreceptor regeneration. Through transient knockdowns of these master regulators, we show that *e2f1*, *e2f2*, *e2f3*, and *aurkb* play critical roles in progenitor cell proliferation. We also show that without affecting proliferation, *prdm1a* is important for the differentiation of progenitor cells into rod photoreceptors. We also show that when differentiation into rods is inhibited, these progenitor cells do not differentiate into other cell types. Overall, this study reveals the transcription factors necessary for rod photoreceptor regeneration in a Zf model with chronic retinal degeneration and regeneration.

## Results

### Rod degeneration and progenitor cell populations in the P23H Zf retina

In previous studies, we have used immunohistochemistry and Single-Cell RNA Sequencing (SC-RNA Seq) analyses on retinas from adult WT and P23H transgenic Zf (expressing Flag-tagged P23H mutant rhodopsin in rods) to characterize how the retinal environment changed during continuous rod photoreceptor degeneration and regeneration (26, 27). The degeneration of rod photoreceptors in the P23H Zf is characterized by a decrease in the number of rod photoreceptors as compared to WT (Figure 1A, 1C; (26)). Furthermore, the rod photoreceptors that are present have much smaller outer segments as compared to WT (Figure 1A, 1C blue arrows; (26)). It should be noted that the Retp1 antibody against the amino terminus of rat rhodopsin labels both zebrafish rods and the green cone outer segments (orange arrows, Figure 1C) due to highly similar amino acid sequences of the epitope recognized. SC-RNA Seq analysis reveals that the P23H retina contains a distinct rod population not found in WT (Figure 1B, 1D; (27)). This cluster contains genes found unique to both retinal progenitor cell and rod photoreceptor populations (Table 1). We believe this cluster to be new rod photoreceptors derived from progenitor cells (Figure 1D).

**Figure 1.**
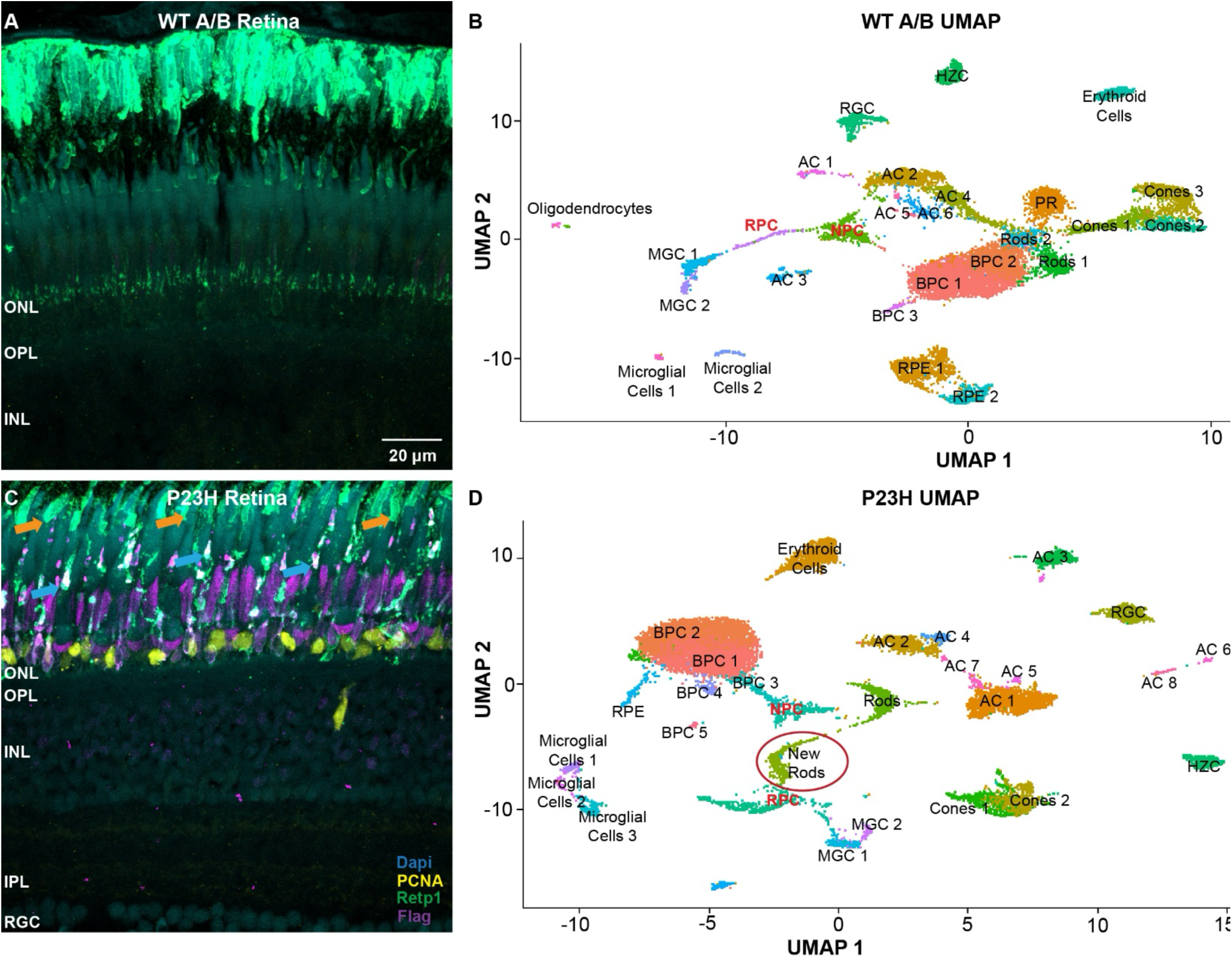
Retinal landscape of WT and P23H retinas. A) Immunostaining for PCNA (yellow), rhodopsin (green), and Flag tag (P23H rhodopsin transgene; magenta) in adult WT retina. B) UMAP projection of the different cell clusters identified through SC-RNA Seq. C) Immunostaining for PCNA (yellow), rhodopsin (green), and Flag tag (magenta) in adult P23H retina. Orange arrows point to off-target labeling of green cone photoreceptor outer segments by Retp1 antibody. Blue arrows point to deformed outer segments of rod photoreceptors. Note that the ONL of the P23H retina is much thinner than WT retina. D) UMAP projection of the different cell clusters identified through SC-RNA Seq. P23H retinas have a unique cluster identified as new rods (circled in red). ONL=outer nuclear layer (photoreceptors), OPL= outer plexiform layer (photoreceptor synapses), INL= inner nuclear layer (horizontal, bipolar, amacrine and Müller cells), IPL= inner plexiform layer (synapses), RGC= retinal ganglion cells, AC= amacrine cells, BPC= bipolar cells, HZC = horizontal cells, RPE = retinal pigmented epithelial cells, MGC= Müller glial cells, RPC= retinal progenitor cells, NPC= neurogenic progenitor cells.

**Table 1.** Genes used to identify new rod cluster.

| Gene | Cell Type | Gene ID | Source |
| --- | --- | --- | --- |
| *crx* | Photoreceptors | ZDB-GENE-010403-1 | The developmental sequence of gene expression within the rod photoreceptor lineage in embryonic zebrafish (74). |
| *otx2* | Photoreceptors | ZDB-GENE-980526-406 | Functional roles of Otx2 transcription factor in postnatal mouse retinal development (77). |
| *pde6ga* | Rod Photoreceptors | ZDB-GENE-050522-144 | Evolution and expression of the phosphodiesterase 6 genes unveils vertebrate novelty to control photosensitivity (89). |
| *pde6gb* | Rod Photoreceptors | ZDB-GENE-030904-1 | Evolution and expression of the phosphodiesterase 6 genes unveils vertebrate novelty to control photosensitivity (89). |
| *fabp7a* | Progenitor Cells | ZDB-GENE-000627-1 | Molecular characterization of retinal stem cells and their niches in adult zebrafish (90). |
| *hmga1a* | Progenitor Cells | ZDB-GENE-010502-1 | Gene regulatory networks controlling vertebrate retinal regeneration (50). |
| *insm1a* | Progenitor Cells | ZDB-GENE-040426-1810 | Insm1a-mediated gene repression is essential for the formation and differentiation of Müller glia-derived progenitors in the injured retina (91). |
| *neurod1* | Progenitor Cells | ZDB-GENE-990415-172 | Overexpressing NeuroD1 reprograms Müller cells into various types of retinal neurons (92). |
| *sox11* | Progenitor Cells | ZDB-GENE-980526-395 | The early retinal progenitor-expressed gene Sox11 regulates the timing of the differentiation of retinal cells (93). |

Consistent with our previous studies, progenitor cells labeled with Proliferating Cell Nuclear Antigen (PCNA) antibody were found predominantly in the ONL in our P23H Zf model (Figure 1C; (28)). SC-RNA Seq analysis revealed 2 progenitor cell clusters in the P23H data that have previously been identified as Neurogenic Progenitor Cells (NPCs) and Retinal Progenitor Cells (RPCs) (Figure 1D; (27)). We performed HiPlex in-situ hybridizations to visually differentiate the two clusters. *Fam60al* was a gene found to be predominantly unique to and highly expressed in both progenitor cell clusters (Figure 2A). *Aurkb* was found to be predominantly unique to and highly expressed only in the RPC cluster (Figure 2B). To differentiate between the clusters visually, RPCs were expected to express both *fam60al* and *aurkb*, while NPCs were expected to express only *fam60al*. HiPlex data showed that cells expressing both *fam60al* and a*urkb* were found predominantly in the ONL, while cells expressing only *fam60al* were predominantly located in the inner nuclear layer (INL) (Figure 2C, D, E). Thus, we identified the PCNA-labeled cell population present in the ONL as the RPC population and the NPC population to be an unlabeled cell class located in the INL.

**Figure 2.**
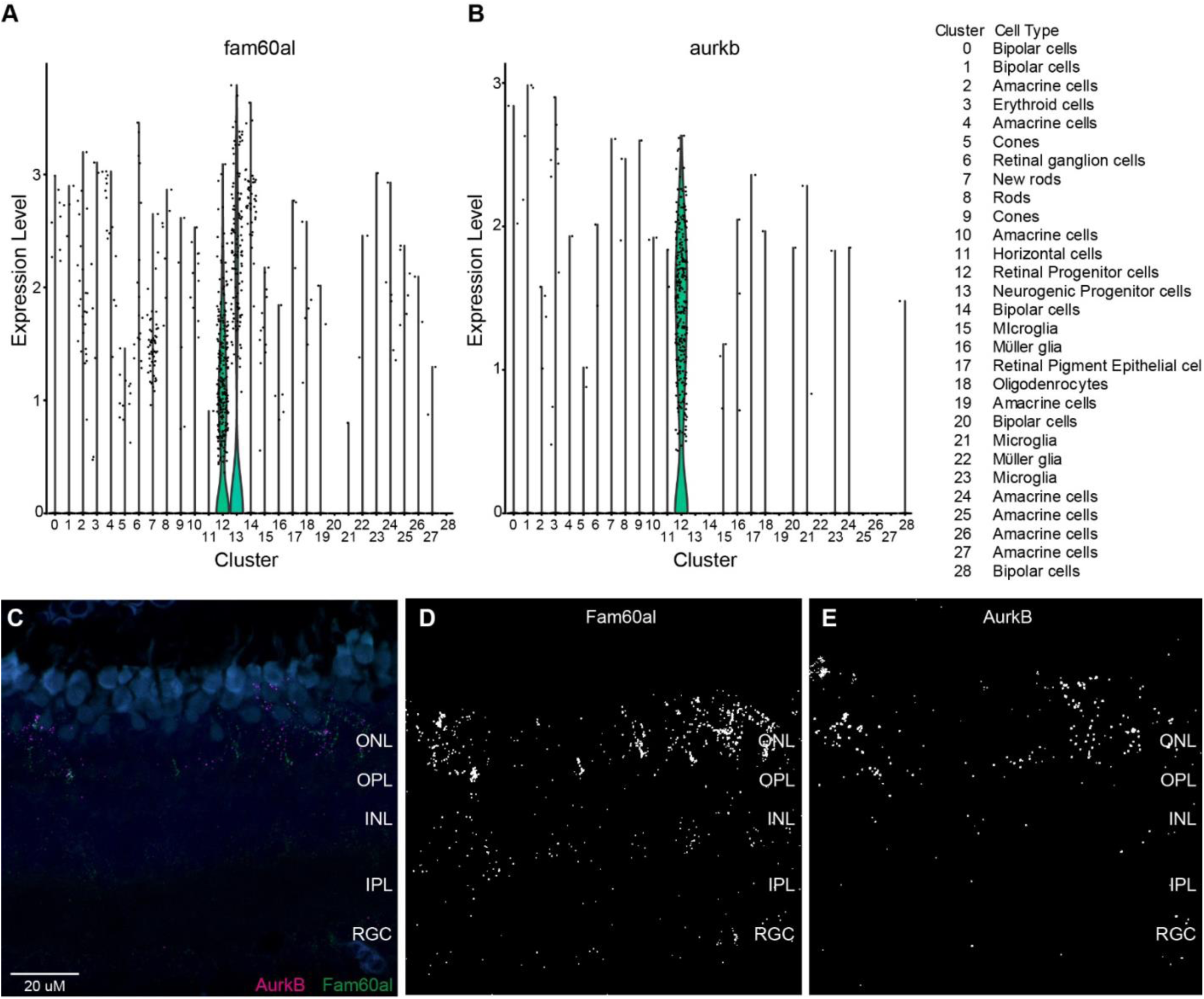
Identification of progenitor cell types in the retina. A) Violin plot of *fam60al*, a gene that is uniquely and highly expressed in both progenitor cell clusters (RPC, cluster 12; NPC, cluster 13). B) Violin plot of *aurkb*, a gene that is uniquely and highly expressed only in the retinal progenitor cell cluster. C) Detection of *fam60al* and *aurkb* transcripts through in-situ hybridization in adult Zf retina. D) Detection of *fam60al* transcripts through in-situ hybridization. *Fam60al* is detected in both the ONL and INL regions of the retina. E) Detection of *aurkb* transcripts through in-situ hybridization. *Aurkb* is detected predominantly in the ONL region of the retina.

### Rapid turnover of progenitor cells and rods in P23H Zf retina

To assess the rate of regeneration and degeneration, we performed a Bromodeoxyuridine (BrdU) pulse-chase experiment. Both WT and P23H Zf were given intraperitoneal injections of BrdU for 3 consecutive days. Zf were then collected 5 hours after the last injection as well as 7 days after the last injection. Figure 3A shows a timeline schematic of the injection and collection timepoints. Wholemounts of retinas were collected and immunostained for rhodopsin (Retp1) and BrdU. WT retinas collected on the last day of injections and imaged at the level of the ONL showed rhodopsin detected in a mosaic pattern (Figure 3B), consistent with modest intracellular labeling in the cell bodies in the ONL (Figure 1A). BrdU labeling was very sparse and irregularly found (Figure 3B), amounting to 30.8 ± 9.4 labeled nuclei per 40x image field (Figure 3H). A 3D projection of the wholemount revealed that BrdU-positive cells were sparsely distributed in both the ONL and INL of the retina (Figure 3D). WT retinas collected 7 days after the last injection showed the same mosaic labeling of rhodopsin with no significant changes to BrdU labeling throughout the retina (28.2 ± 8.8 nuclei/field; p = 1.0; Figure 3C, H). P23H Zf retinas collected on the 3^rd^ day of injections did not have a regular mosaic labeling pattern for rhodopsin because rhodopsin was delocalized over the entire rod photoreceptor and was visible in somata imaged in the ONL (Figure 3E). Furthermore, P23H retinas had a significantly higher number of BrdU-positive cells when compared to WT (684.2 ± 27.2 nuclei/field; p < 0.001; Figure 3H). A 3D projection of the wholemount revealed that BrdU-positive cells were predominantly found in the ONL, with very few in the INL (Figure 3G), suggesting that the rapidly proliferating cell population is the one we identified as RPCs. P23H retinas collected 7 days after the last injection still did not have a mosaic pattern of rhodopsin labeling in the ONL and the number of BrdU positive cells (299.0 ± 26.4 nuclei/field) was still significantly higher than WT (28.2 ± 8.8; p < 0.0001; Figure 3F, H). Furthermore, a portion of BrdU-positive cells were also rhodopsin positive, indicating that the RPCs had differentiated into rod photoreceptors (Figure 3F). However, the number of BrdU-positive cells after 7 days (299.0 ± 26.4) was significantly reduced compared to the number found in P23H retinas collected on the 3^rd^ day of injections (684.2 ± 27.2; p < 0.001; Figure 3H). Thus, 56% of BrdU-labeled cells were lost in 7 days in the P23H Zf model, suggesting that the half-life of a regenerated rod photoreceptor is less than a week.

**Figure 3.**
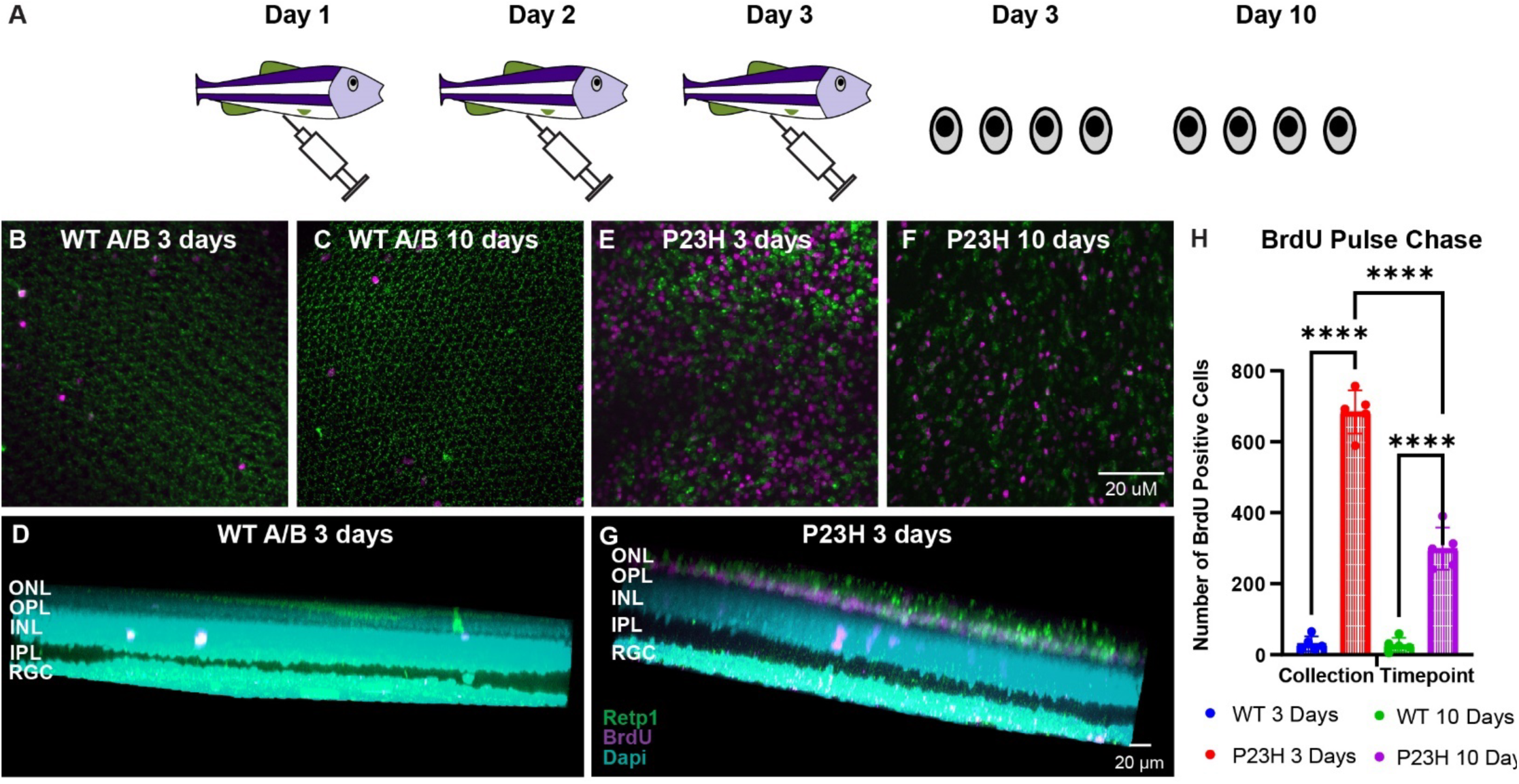
BrdU pulse-chase in adult Zf retina. A) Timeline schematic of experimental procedures. B) Rhodopsin and BrdU labeling in WT A/B retina collected on the 3^rd^ day of BrdU injections. C) Rhodopsin and BrdU labeling in WT A/B retina collected 7 days after the last BrdU injection. D) 3-D image of WT A/B retina collected on the 3^rd^ day of BrdU injections. E) Rhodopsin and BrdU labeling in P23H retina collected on the 3^rd^ day of BrdU injections. F) Rhodopsin and BrdU labeling in P23H retina 7 days after the last BrdU injection. G) 3-D image of P23H retina collected on the 3^rd^ of BrdU injections. H) Quantification of the number of BrdU positive cells at each timepoint. Data points represent individual animals. **** p < 0.0001; n = 5 animals/condition.

### Prediction of transcriptional programs required for rod photoreceptor regeneration

The high turnover rate of regenerated rod photoreceptors in the P23H transgenic model was advantageous for the analysis of the transcriptional mechanisms of rod regeneration, as cells at all stages of differentiation were present in our SC-RNA Seq datasets. This allowed us to evaluate trajectories of gene expression changes from potential progenitor cell populations to mature rods within the same dataset. To identify the transcription factors responsible for rod photoreceptor regeneration in the P23H Zf model, SC-RNA Seq data were first run through Monocle3. Monocle3 uses a machine learning algorithm to identify the changes in gene expression needed to go from one cell type (cluster) to another cell type (29, 30). Trajectories are plotted through expression score analyses as well as pseudotime analyses. Predicted trajectories were selected based on having a high expression score and a low pseudotime. When WT data were processed, one trajectory was observed to predict the differentiation of RPCs and NPCs into amacrine cells, bipolar cells, and rod photoreceptors (Figure 4A). When P23H data were processed, three trajectories were predicted. The first trajectory predicted a similar differentiation of RPCs and NPCs into amacrine cells, bipolar cells and rods (specifically new rods) (Figure 4B). The second trajectory predicted the differentiation of RPCs (and some NPCs) into new rods (Figure 4C). The third trajectory predicted the maturation of new rods (Figure 4D). The network of genes involved in each of the trajectories found from the P23H data were then run through DrivAER analysis to identify potential master regulators of these networks (31). DrivAER analysis predicted 49 transcription factor master regulators of the proliferation of progenitor cells; however, *e2f1*, *e2f2*, and *e2f3* had the greatest number of trajectory gene hits (Figure 5A; Supplementary Table 1). Differentiation of progenitor cells into rod photoreceptors had 16 predicted master regulators, with *prdm1a* having the greatest number of trajectory gene hits (Figure 5B; Supplementary Table 1). Maturation of rod photoreceptors only had 2 predicted transcription factor master regulators (Figure 5C; Supplementary Table 1). It was predicted that e2*f1, e2f2, and e2f3* were responsible for the proliferation of progenitor cells, *prdm1a* was responsible for the differentiation of progenitor cells into rod photoreceptors, and *sp1* was responsible for the maturation of rod photoreceptors (Figure 5D). The list of genes from the predicted trajectories regulated by each transcription factor is provided in Supplementary Figures 1-3. The remainder of this study will explore the proliferation and differentiation aspects of rod photoreceptor regeneration.

**Figure 4.**
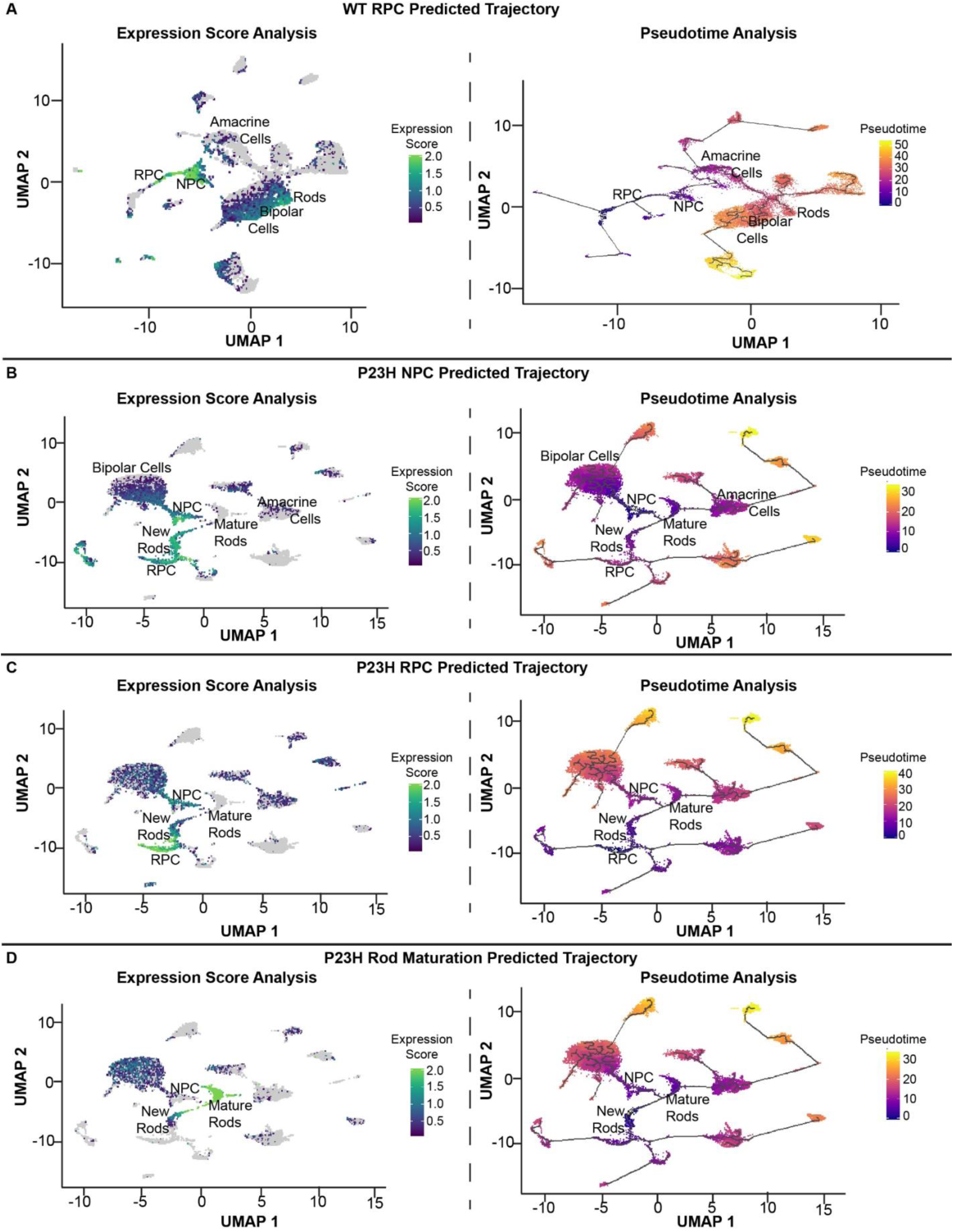
Trajectory analysis of WT and P23H SC-RNA seq data. A) Expression score analysis and pseudotime analysis of the progenitor cell populations found in the WT data. B) Expression score analysis and pseudotime analysis of the neurogenic progenitor cell cluster in the P23H data. C) Expression score analysis and pseudotime analysis of the retinal progenitor cell cluster in the P23H data. D) Expression score analysis and pseudotime analysis of rod photoreceptor maturation. The higher the expression score and lower the pseudotime, the more likely the predicted trajectory will happen.

**Figure 5.**
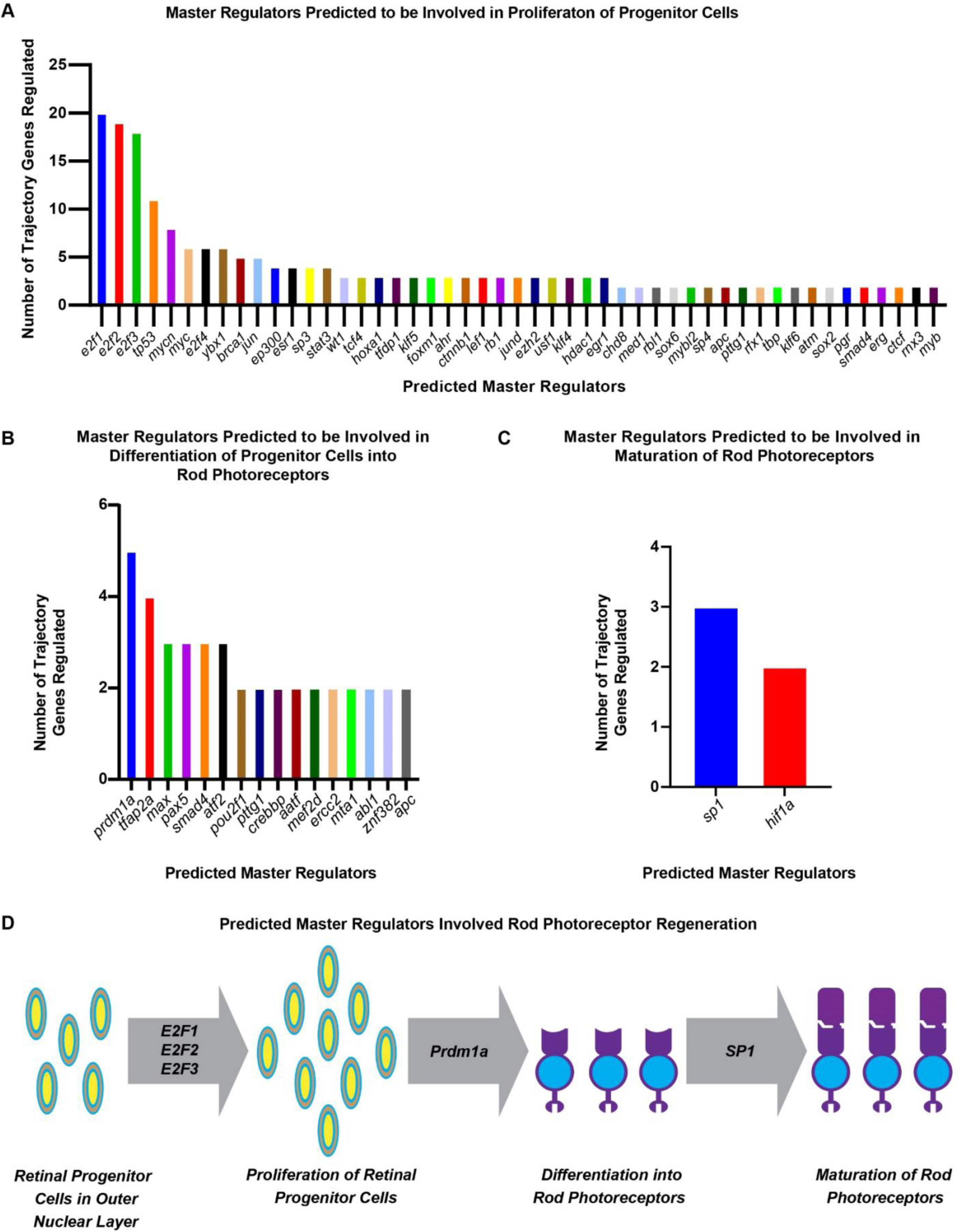
DrivAER prediction of master regulators involved in rod photoreceptor regeneration. A) Transcription factors regulating genes involved in proliferation of RPCs. B) Transcription factors regulating genes involved in differentiation of RPCs into new rods. C) Transcription factors regulating genes involved in maturation of new rods. D) *E2f1, e2f2,* and *e2f3* are predicted to play an important role in the proliferation of progenitor cells in the ONL. *Prdm1a* is predicted to play an important role in the differentiation of progenitor cells into rod photoreceptors. *Sp1* is predicted to play an important role in the maturation of newly formed rod photoreceptors.

### Proliferation knockdown in P23H Zf retina

To test whether master control genes identified in the DrivAER analysis are central to the processes they were associated with, we performed targeted knock-downs of the top hits associated with each process in adult P23H zebrafish retina. Proliferation of the RPC population is the first prerequisite for regeneration of rods, and members of the *e2f* family of transcription factors were identified as the top regulators of this process. To test the effects of *e2f1, e2f2,* and *e2f3* on proliferation, translation-blocking Vivo-morpholino oligonucleotides were generated to target each of the transcription factors (Table 2). Adult P23H Zf were intravitreally injected with a cocktail of the *e2f1, e2f2,* and *e2f3* Vivo-morpholinos or a control Vivo-morpholino, followed by intraperitoneal injections of BrdU on the following two days (Figure 6A). Once Zf retinas were collected and processed, tissue slices were immunostained and quantified for BrdU labeling. 5 hr after the last BrdU injection, control retina slices had 15.0 ± 3.1 BrdU positive cells per 40x image field (Figure 6B, H). Tissue slices collected from retinas injected with a cocktail of *e2f1, e2f2,* and *e2f3* Vivo-morpholinos had a significant reduction in BrdU positive cells when compared to the controls (p = 0.0439) with 4.2 ± 1.9 BrdU positive cells per image field (Figure 6C, H). *Aurkb* is known to play an important role in the G2/M phase of the cell cycle (32) and has been found to regulate regeneration of spinal motor neurons in developing Zf (32–34). It was unique to RPCs in our P23H SC data and hence we have generated a translation blocking Vivo-morpholino to target *aurkb* (Table 2). Adult P23H Zf were intravitreally injected with the *aurkb* Vivo-morpholino followed by intraperitoneal injections of BrdU at similar time points as the *e2f* and control conditions. Tissue slices collected from retina injected with the *aurkb* Vivo-morpholino had a significant reduction in BrdU positive cells when compared to the control (p = 0.0239) with 2.5 ± 2.1 BrdU positive cells per image field (Figure 6D, H). There was no significant difference between BrdU positive cell count between *aurkb* and *e2fs* Vivo-morpholinos condition (p = 0.8689), indicating that both are essential for RPC proliferation.

**Figure 6.**
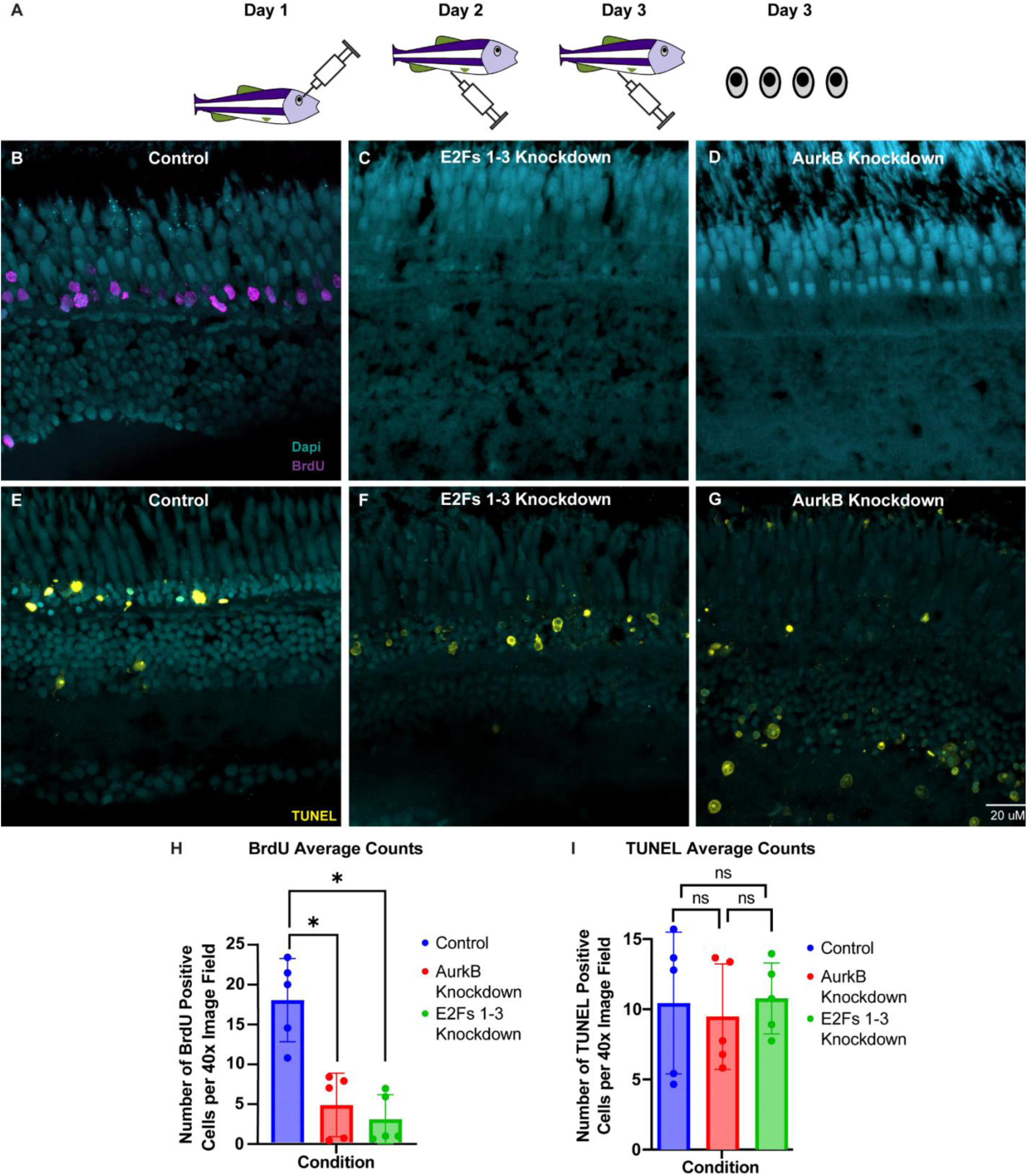
Effects of *e2f’s 1-3* and *aurkb* knockdowns on RPC proliferation. A) Timeline schematic of experimental procedures. B) BrdU labelling of control retina. C) BrdU labelling of retina injected with a cocktail of *e2f1, e2f2,* and *e2f3* Vivo-morpholinos. D) BrdU labelling of retina injected with the *aurkb* Vivo-morpholino. E) TUNEL staining of control retina. F) TUNEL staining of retina injected with a cocktail of *e2f1*, *e2f2*, and *e2f3* Vivo-morpholinos. G) TUNEL staining of retina injected with the *aurkb* Vivo-morpholino. H) Quantification of the number of BrdU positive cells found within each treatment. Both *e2f’s 1-3* and *aurkb* knockdown significantly reduced the number of BrdU-positive proliferating progenitor cells. I) Quantification of the number of TUNEL positive cells found within the ONL in each treatment. Neither *e2f’s 1-3* nor *aurkb* knockdown altered the number of TUNEL-positive cells in the ONL. Data points represent individual animals. * p < 0.05; n = 5 animals/condition.

**Table 2.**
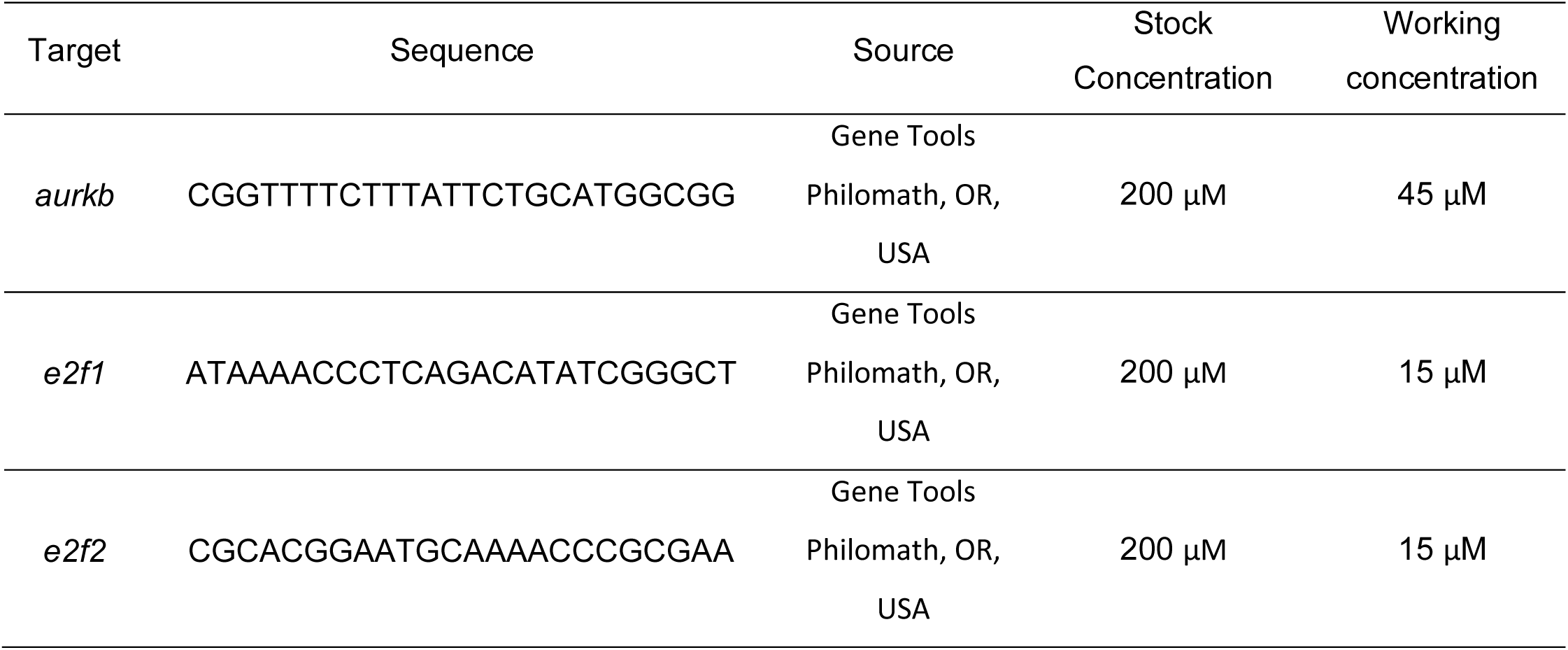
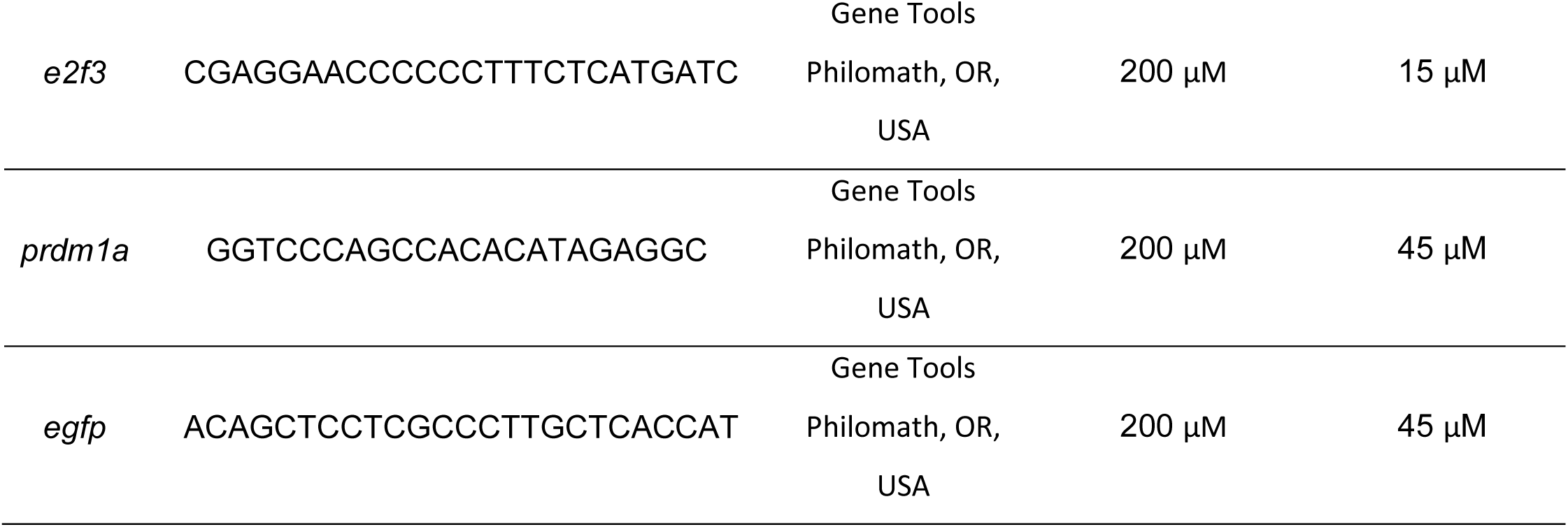
Vivo-Morpholinos for targeted gene knockdowns.

While reduced proliferation of progenitor cells is the most likely explanation for the reduced incorporation of BrdU, an alternative explanation could be that knockdown or *e2fs 1-3* or *aurkb* triggered apoptosis in retinal progenitor cells or newly formed rods, reducing the number of BrdU-labeled cells. To investigate this hypothesis, we performed a TUNEL staining assay to identify cells undergoing apoptosis. Control retina slices had 12.1 ± 1.3 TUNEL positive cells in the ONL per 40x image field (Figure 6E, I). Tissue slices collected from retinas injected with a cocktail of *e2f1*, *e2f2*, and *e2f3* Vivo-morpholinos had no significant reduction in TUNEL positive cells in the ONL when compared to the controls (p = 0.7266) with 9.7 ±3.3 TUNEL positive cells per 40x image field (Figure 6F, I). While retinas injected with the *aurkb* Vivo-morpholino did display additional TUNEL labeling of unidentified cells in the INL and inner plexiform layer (IPL; Figure 6G), there was no significant change in TUNEL positive cells in the ONL when compared to the controls (p = 0.7266) with 9.7 ± 3.3 TUNEL positive cells per image field (Figure 6G, I). Furthermore, there was no significant difference between TUNEL positive cell count in the ONL between *aurkb* and *e2fs* Vivo-morpholinos condition (p = 0.8563). The decrease in proliferation without an increase in cell death indicates that both *e2fs* and *aurkb* are essential for the proliferation step of regeneration.

To investigate the global effects of *e2fs* and *aurkb* knockdown, retinas undergoing the same experimental paradigm as described above were collected and subjected to liquid chromatography-mass spec (LC-MS-MS) total proteome analysis. When compared to the control retina, retinas with *e2f1*, *e2f2*, and *e2f3* knockdowns had on average a 1.17 log2 fold decrease in expression of the targeted proteins (Supplementary Table 2). Furthermore, expression of proteins known to be directly targeted by the *e2fs*, such as *rrm1*, *ccnyl1,* and *ddx55* were significantly decreased in the knockdown conditions. Biological processes including ncRNA processing, rRNA processing, retina development processes, and photoreceptor cell development processes were downregulated (Figure 7A; Table 3). Conversely, mRNA processing, oxoacid metabolic processes and phototransduction through cone opsins were all upregulated after *e2fs1-3* knockdown (Figure 7A; Table 4). In the *aurkb* knockdown condition, *aurkb* expression was reduced on average by 1.34 log2 fold when compared to the control (Supplementary Table 2). Genes downstream of *aurkb* that are involved in cytokinesis, such as *igf2bp3* and *dock11*, were downregulated (Supplementary Table 2). Biological processes such as histone modification, positive regulation of transcription, covalent chromatin modification and visual perception were downregulated (Figure 7B; Table 5). Conversely, biological processes involved in gene silencing, establishment of mitotic spindle localization and posttranscriptional regulation of gene expression were upregulated (Figure 7B; Table 6).

**Figure 7.**
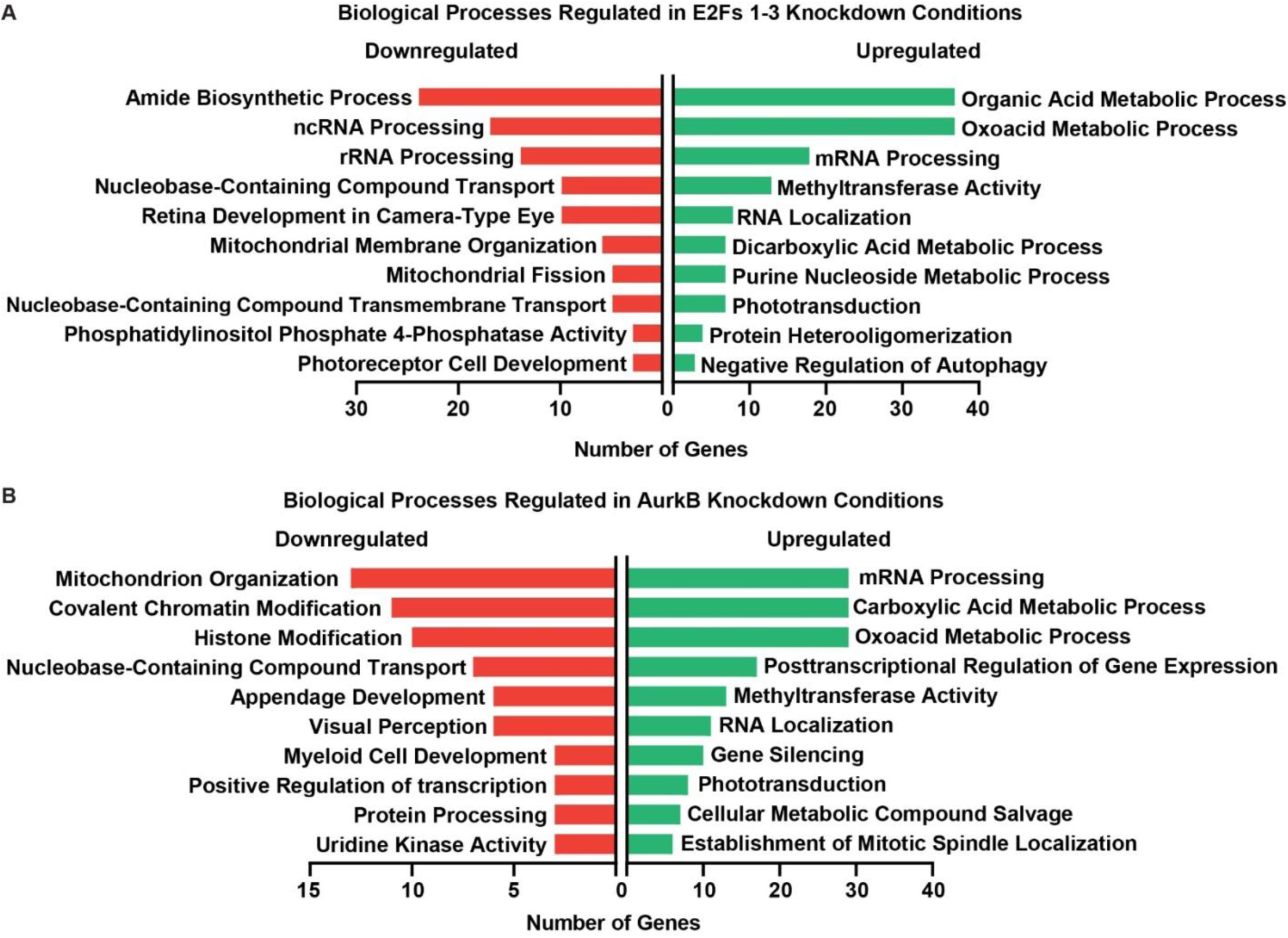
Biological processes affected by Vivo-morpholino knockdown of regulators of RPC proliferation. A) Biological processes downregulated (red; left) and upregulated (green; right) after knockdown of E2Fs 1-3. B) Biological processes downregulated (red; left) and upregulated (green; right) after knockdown of AurkB. Data are derived from two replicate samples per condition.

**Table 3.**
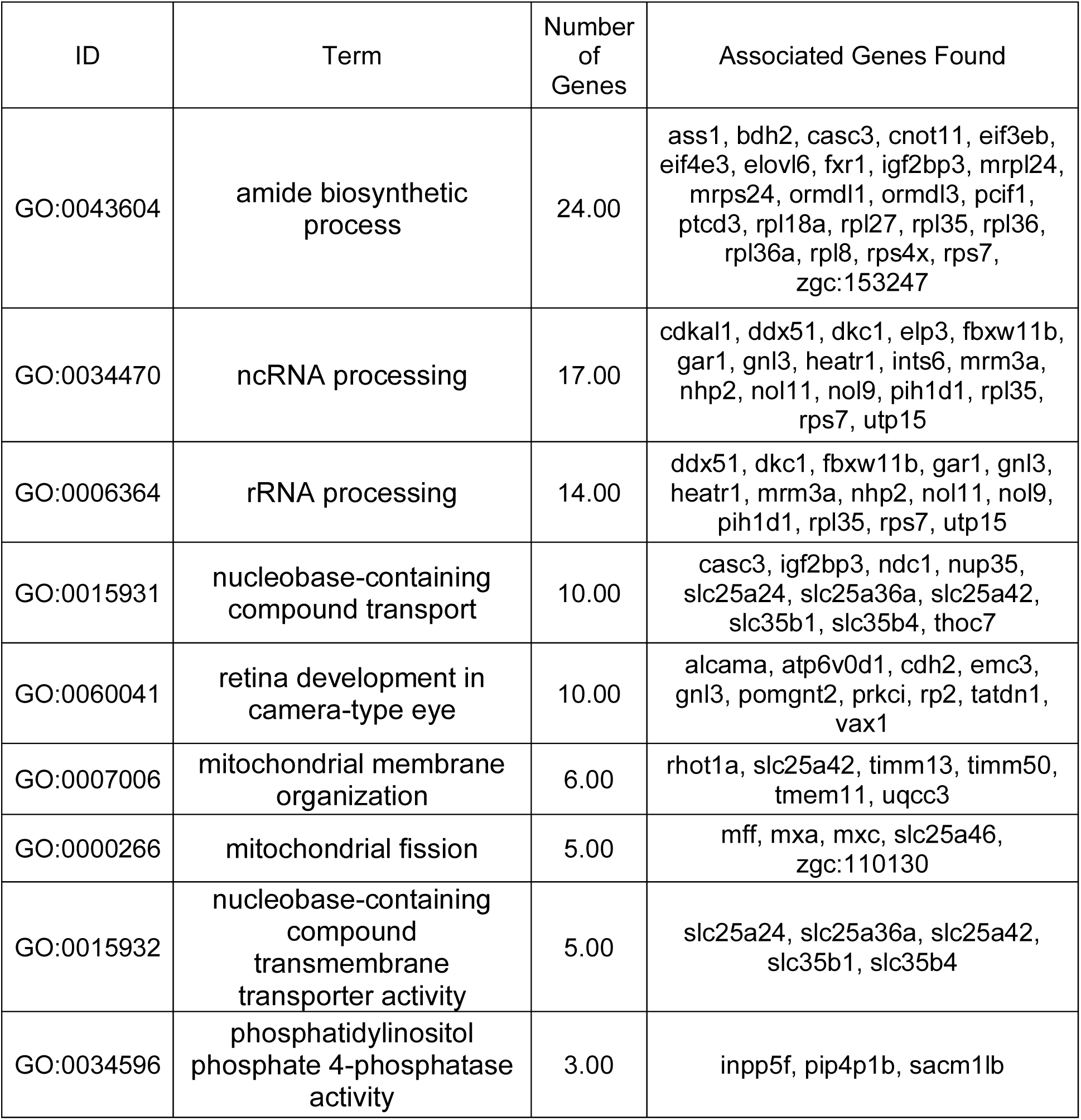
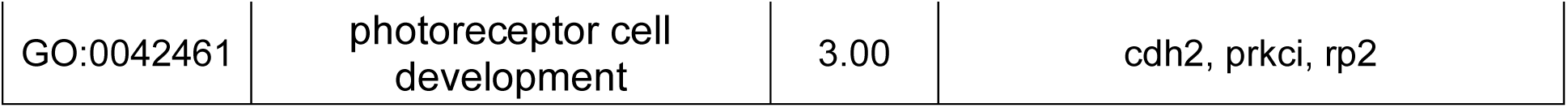
Biological processes downregulated by *e2f1s1-3* knockdown.

**Table 4.**
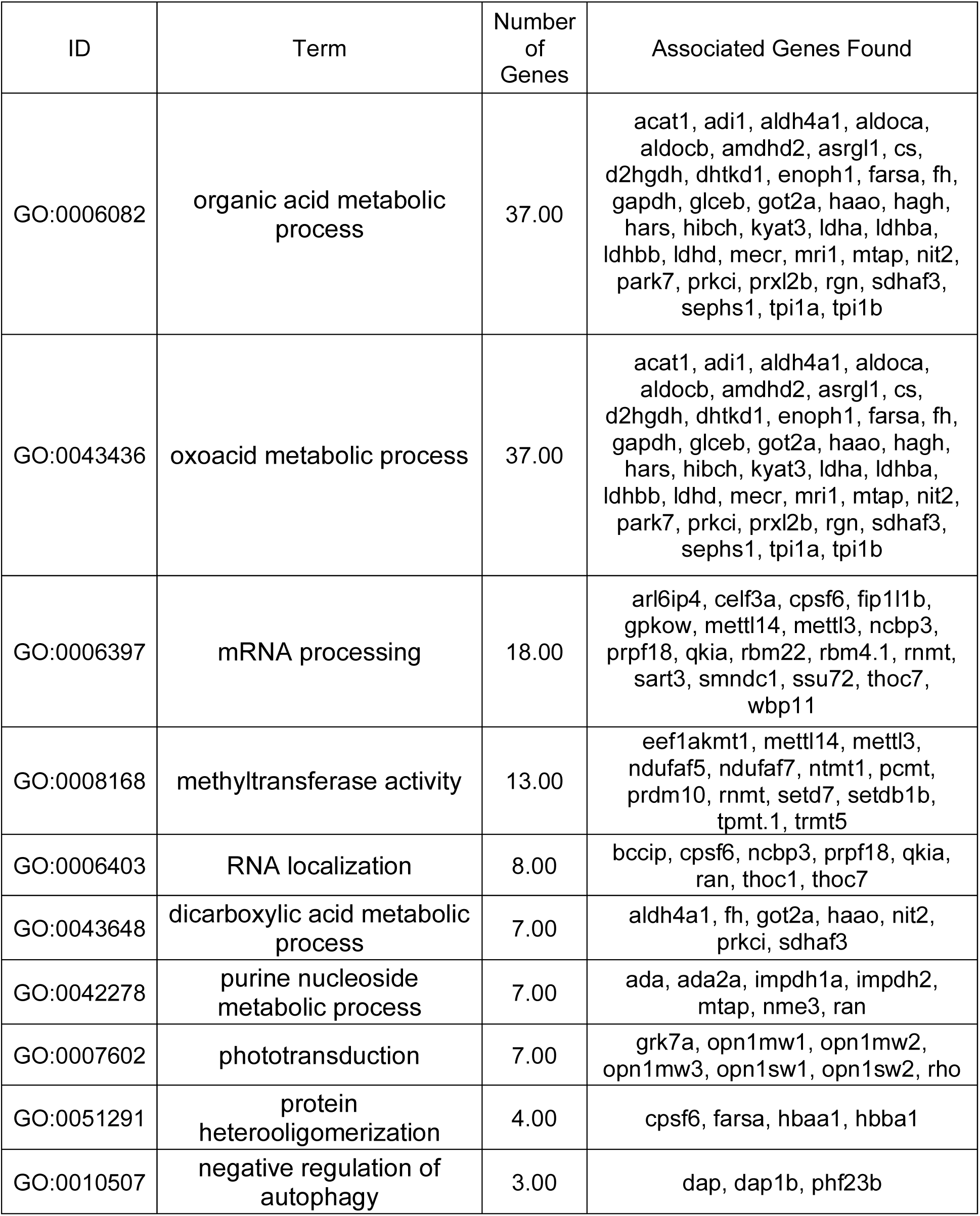
Biological processes upregulated by *e2fs1-3* knockdown.

**Table 5.** Biological processes downregulated by *aurkb* knockdown.

| ID | Term | Number of Genes | Associated Genes Found |
| --- | --- | --- | --- |
| GO:0007005 | mitochondrion organization | 13.00 | atp23, bcs1l, cisd2, immp2l, mtfr1l, mxa, mxc, rhot1a, sdhaf2, slc25a42, timm13, uqcc3, zgc:110130 |
| GO:0016569 | covalent chromatin modification | 11.00 | atf7ip, eed, kat2b, paf1, phf2, setd3, setdb1a, suz12b, tada2b, trrap, waca |
| GO:0016570 | histone modification | 10.00 | eed, kat2b, paf1, phf2, setd3, setdb1a, suz12b, tada2b, trrap, waca |
| GO:0015931 | nucleobase-containing compound transport | 7.00 | casc3, igf2bp3, ndc1, slc25a42, slc35b1, slc35b4, thoc7 |
| GO:0048736 | appendage development | 6.00 | fbln1, fibinb, kat2b, med12, pi4k2b, trim33 |
| GO:0007601 | visual perception | 6.00 | lamc1, opn1mw4, opn4.1, pcdh19, rho, vsx1 |
| GO:0061515 | myeloid cell development | 3.00 | adnpa, bdh2, tpma |
| GO:0045943 | positive regulation of transcription by RNA polymerase I | 3.00 | heatr1, nol11, phf2 |
| GO:0016485 | protein processing | 3.00 | atp23, immp2l, spcs2 |
| GO:0004849 | uridine kinase activity | 3.00 | uck1, uck2a, uck2b |

**Table 6.**
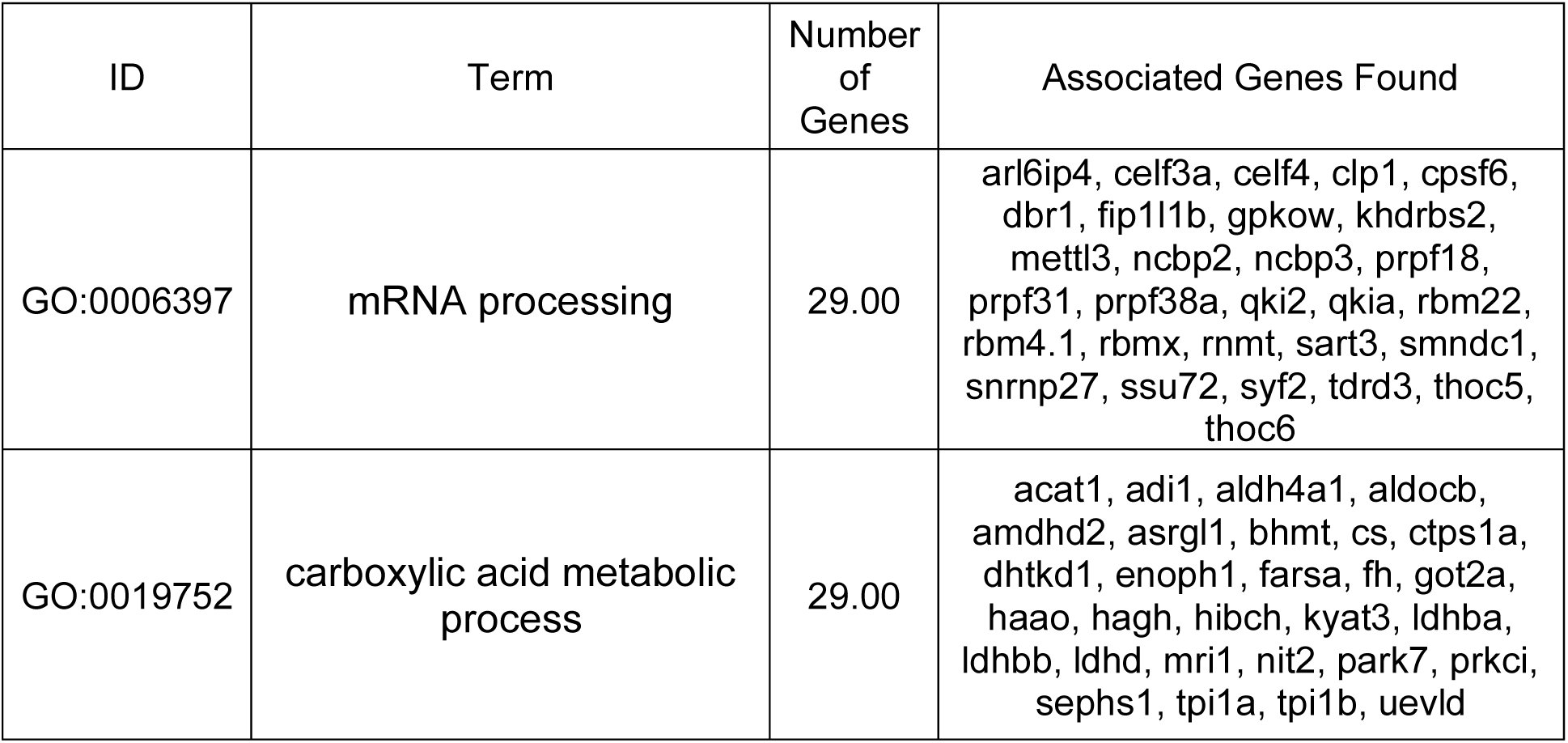
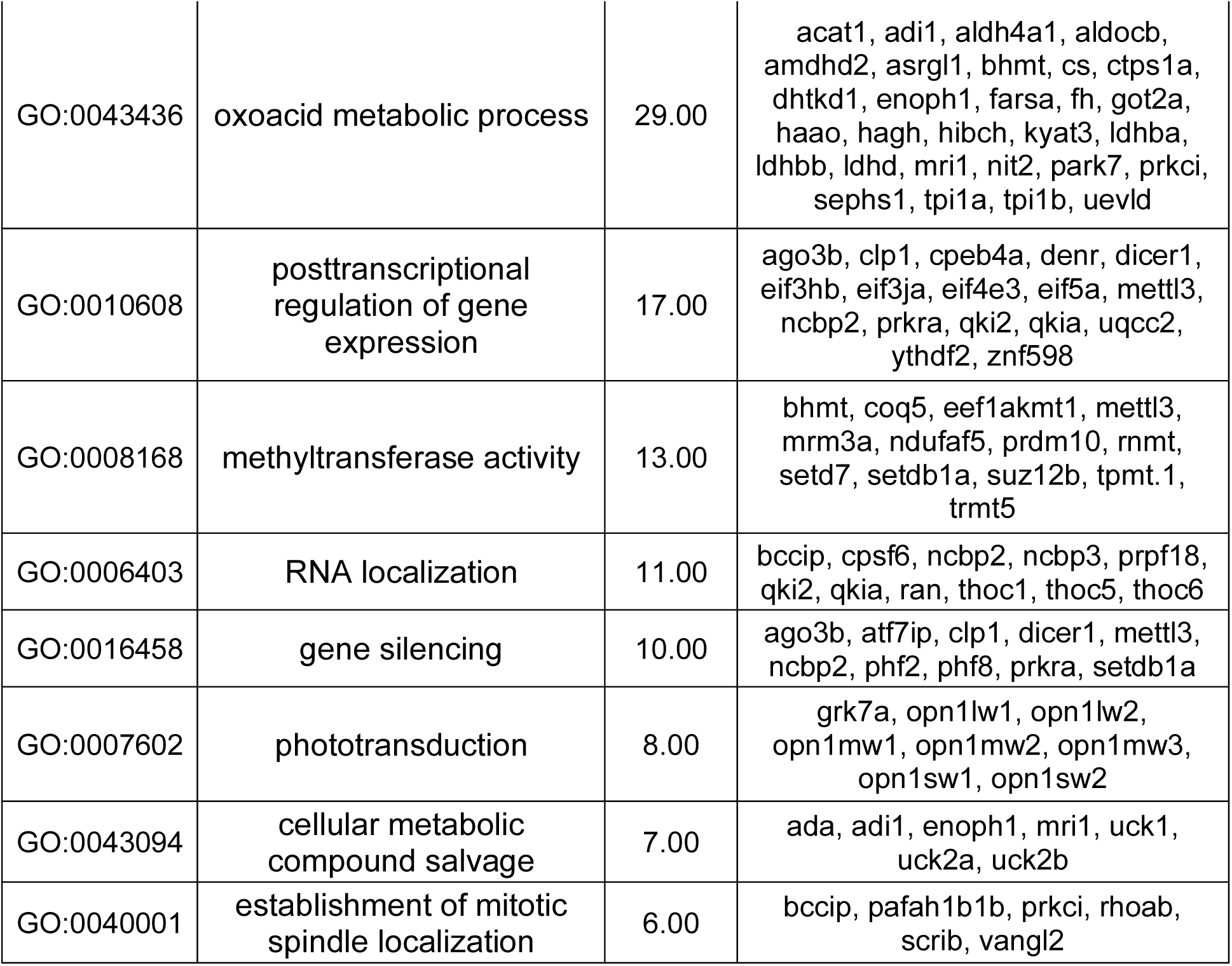
Biological processes upregulated by *aurkb* knockdown.

### Differentiation knockdown in P23H Zf retina

*Prdm1a* was identified as a top transcription factor driving the differentiation of RPCs into rod photoreceptors. To test the effects of *prdm1a*, a translation-blocking Vivo-morpholino was generated to target the transcription factor (Table 2). We first tested whether *prdm1a* knockdown affected the proliferation of progenitor cells. Adult P23H Zf were intravitreally injected with either the *prdm1a* Vivo-morpholino or control Vivo-morpholino, followed by intraperitoneal injections of BrdU at 24 and 48 hpii (Figure 8A). At 53 hpii, BrdU labeling was not significantly reduced compared to the control (p = 0.5944)(Figure 8B-D), while rhodopsin expression was significantly reduced (p = 0.0011)(Figure 8C, D). Because *prdm1a* knockdown did not alter BrdU incorporation into RPCs, we conclude that its effects are independent of the proliferation of progenitor cells.

**Figure 8.**
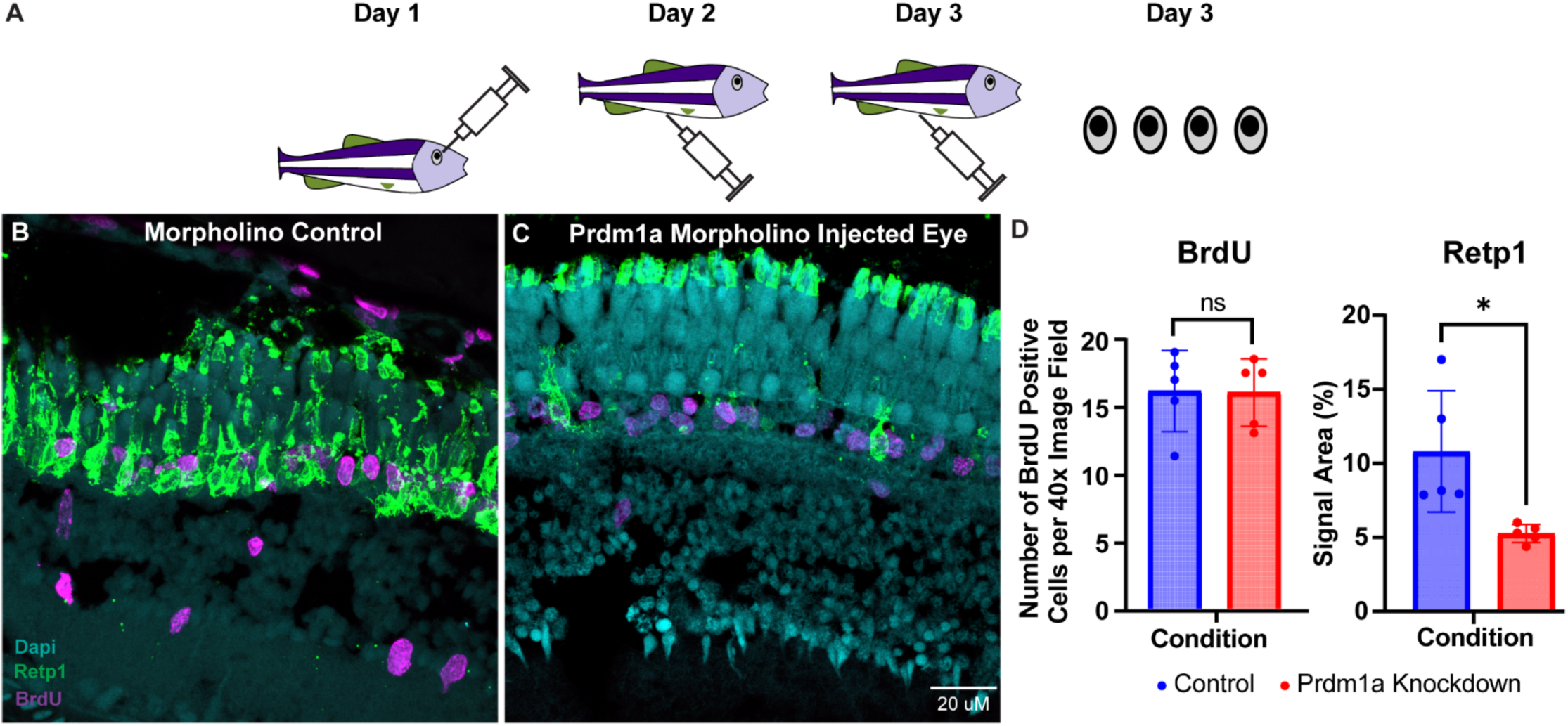
Effects of *prdm1a* knockdown on RPC proliferation and rhodopsin expression in adult P23H retina. A) Timeline schematic of experimental procedures. B) Rhodopsin expression and BrdU labelling of control retina. C) Rhodopsin expression and BrdU labelling of retina injected with the *prdm1a* Vivo-morpholino. D) Quantification of BrdU labelling and rhodopsin expression for each treatment. While the number of BrdU-positive proliferating progenitor cells did not change, *prdm1a* knockdown reduced rhodopsin expression in the ONL. ** p < 0.05; n = 5 animals/condition.

In order to test whether differentiation is affected by *prdm1a* inhibition, adult P23H Zf were given intraperitoneal injections of BrdU first, followed by intravitreal injections of either the *prdm1a* Vivo-morpholino or control Vivo-morpholino. Zebrafish were maintained for 4 days to allow for differentiation of progenitor cells into rod photoreceptors (Figure 9A). Once Zf were collected and processed, tissue slices were immunostained and quantified for BrdU labeling and rhodopsin (Retp1) expression. Rhodopsin expression was significantly lower in tissue slices from the *prdm1a* Vivo-morpholino when compared to the control (p = 0.0314; Figure 9B, C). Surprisingly, the knockdown of *prdm1a* resulted in a significantly higher number of rhodopsin-positive cells in the INL (p = 0.0293; Figure 9C, D). Tissue slices from the control Vivo-morpholino-injected fish had 14.9 ± 1.9 BrdU positive cells present per 40x image field of retina and were not significantly different from tissue slices from the *prdm1a* Vivo-morpholino-injected fish, which had 14.5 ± 1.1 BrdU positive cells present per 40x image field of retina (p = 0.8546; Figure 9B’, C’, E). The number of BrdU-positive cells that colocalized with rhodopsin expression was then used as a measure of progenitor cells differentiating into new rods. In the control group, 94.0 ± 1.2 % of all BrdU-positive cells in the ONL colocalized with rhodopsin expression (Figure 9B’’, F). In the *prdm1a* knockdown group, 62.3 ± 1.8 % of all BrdU positive cells in the ONL colocalized with rhodopsin expression (Figure 9C’’, F). *Prdm1a* knockdown significantly reduced the number of progenitor cells that differentiated into rod photoreceptors (p = 0.0001; Figure 9F). Overall, *prdm1a* knockdown reduced the differentiation of progenitor cells into rod photoreceptors without preventing RPC proliferation.

**Figure 9.**
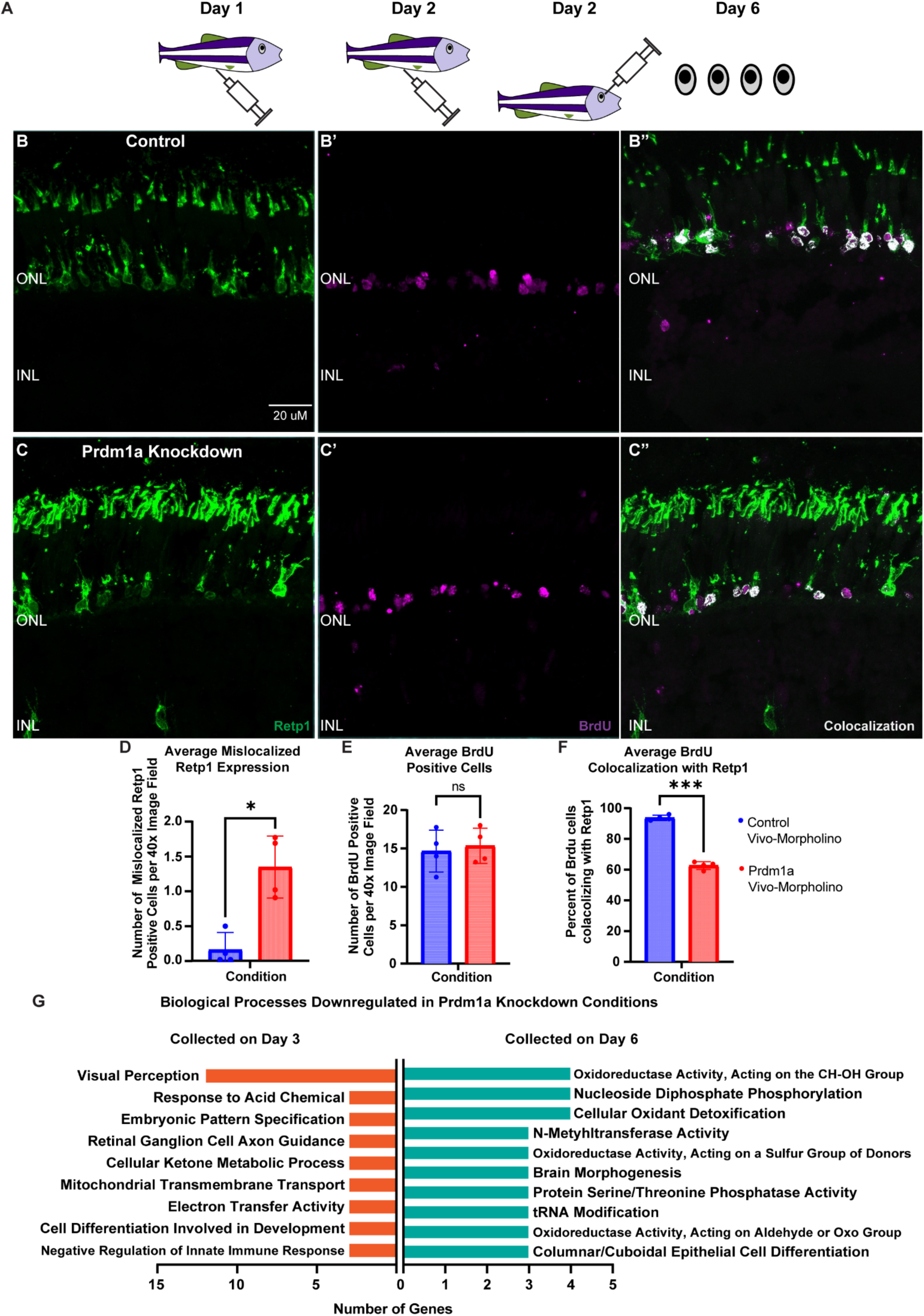
Colocalization of cells that are BrdU-positive and express rhodopsin. A) Timeline schematic of experimental procedures. B) Rhodopsin expression in a control P23H retina. B’) BrdU labelling in a control retina. B’’) Colocalization of cells that are BrdU-positive and express rhodopsin. C) Rhodopsin expression in a P23H retina after *prdm1a* knockdown. C’) BrdU labelling in a P23H retina after *prdm1a* knockdown. C’’) Colocalization of cells that are BrdU positive and express rhodopsin. D) The number of mislocalized cells found in the INL that expressed rhodopsin increased significantly after *prdm1a* knockdown. E) The number of BrdU-positive cells did not change after *prdm1a* knockdown. F) The fraction of BrdU-positive cells that also expressed rhodopsin was significantly reduced after *prdm1a* knockdown. *** p < 0.001, * p < 0.05; n = 4 animals/condition. G) Biological processes downregulated after Prdm1a knockdown at Day 3 (orange;left) and Day 6 (cyan;right) of collection. Data are derived from two replicate samples per condition.

Retinas undergoing the same experimental paradigm as described above were collected and subjected to LC-MS-MS total proteome analysis. When compared to control conditions *prdm1a* knockdowns showed on average a 1.38 log2 fold reduction of targeted protein (Supplementary Table 2). *Prdm1a* knockdowns also showed a significant downregulation of *rho*, a gene uniquely expressed in rod photoreceptors. When tissue samples were collected at 53 hpii, biological processes such as visual perception, cell differentiation and negative regulation of innate immune response were downregulated (Figure 9G; Table 7). When tissue samples were collected 4 days after administration of the *prdm1a* Vivo-morpholino, biological processes such as brain morphogenesis, columnar/cuboidal epithelial cell differentiation and cellular oxidant detoxification were downregulated (Figure 8E; Table 8). The downregulation of biological processes involved in differentiation and neuronal morphogenesis coincide with the reduction of progenitor cell differentiation seen through immunohistochemistry.

**Table 7.** Biological processes downregulated by *prdm1a* knockdown (collection at 53 hpii).

| ID | Term | Number of Genes | Associated Genes Found |
| --- | --- | --- | --- |
| GO:0007601 | visual perception | 12.00 | grk7b, nr2f5, opn1lw1, opn1lw2, opn1mw2, opn1mw3, opn1mw4, opn1sw1, opn1sw2, pcdh19, rho, vsx1 |
| GO:0001101 | response to acid chemical | 3.00 | ache, meak7, nr2f5 |
| GO:0009880 | embryonic pattern specification | 3.00 | cactin, ndr2, prrc1 |
| GO:0031290 | retinal ganglion cell axon guidance | 3.00 | mycbp2, ndr2, rbpms2a |
| GO:0042180 | cellular ketone metabolic process | 3.00 | aldh8a1, coq5, coq8b |
| GO:1990542 | mitochondrial transmembrane transport | 3.00 | chchd4a, slc25a36a, slc25a38b |
| GO:0009055 | electron transfer activity | 3.00 | glrx5, ndr2, ndufa4 |
| GO:0061005 | cell differentiation involved in kidney development | 3.00 | coq8b, prkci, thsd7aa |
| GO:0045824 | negative regulation of innate immune response | 3.00 | b2m, pias4a, samhd1 |

**Table 8.**
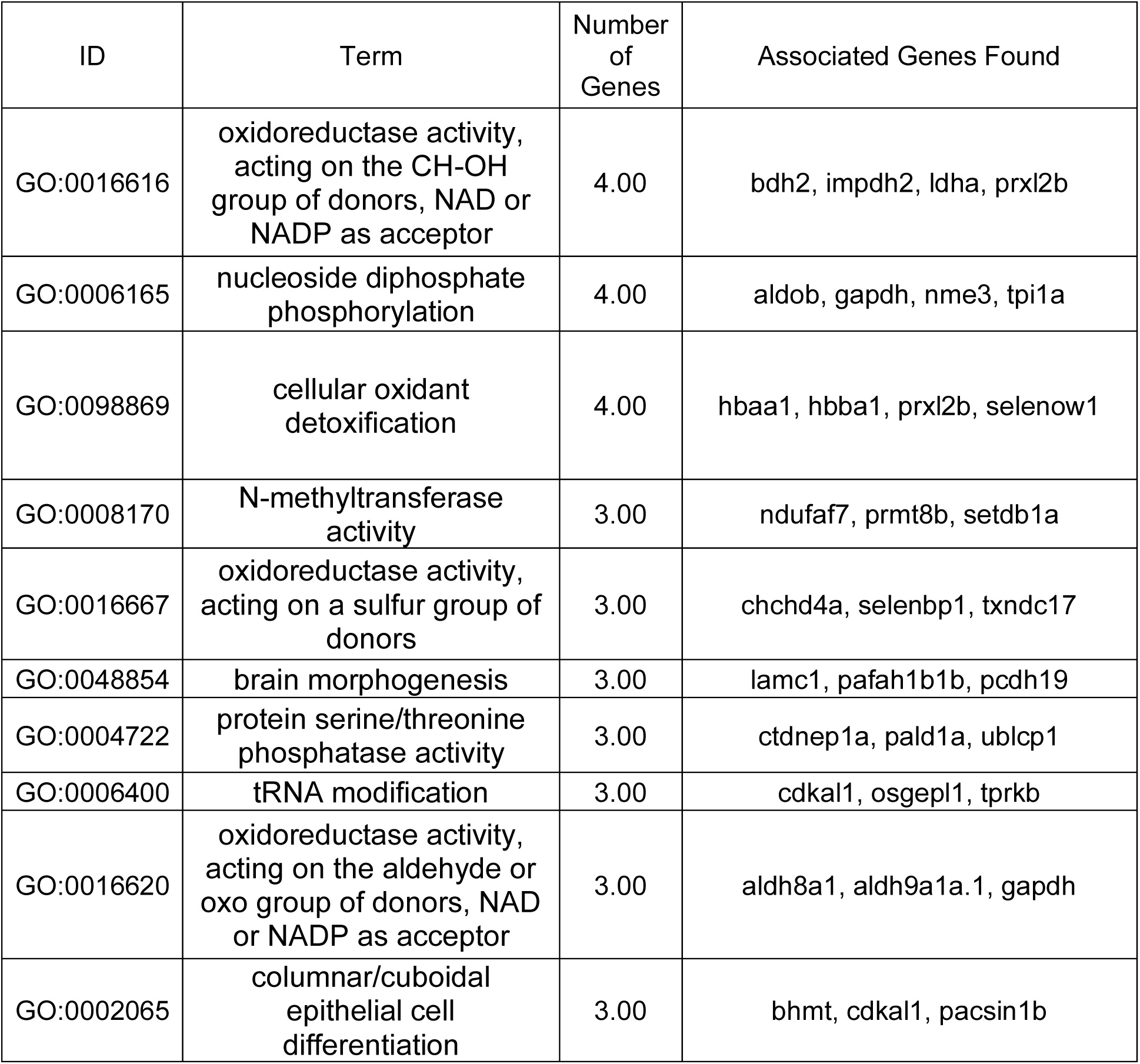
Biological processes downregulated by *prdm1a* knockdown (collection on Day 6).

Because *prdm1a* knockdown significantly reduced the number of BrdU-positive cells that differentiated into rod photoreceptors, we wanted to investigate whether or not those BrdU-positive cells were differentiating into other cell types. Tissue slices from the control group and *prdm1a* knockdown group were stained for BrdU, Pkcα (a protein predominantly expressed in bipolar cells), and ZPR-1 (an antibody that specifically labels cone arrestin in red and green cones) (Figure 10A, B, C). BrdU-positive cells did not label for either Pkcα or ZPR-1 (Figure 10D). Similarly, using blue opsin and UV opsin antibodies to label blue and UV cones, we found no BrdU-positive nuclei associated with either cone subtype (Figure 10E, F). It appears that *prdm1a* knockdown inhibits the differentiation of progenitor cells without pushing progenitor cells to other cell fates such as bipolar cells or cones.

**Figure 10.**
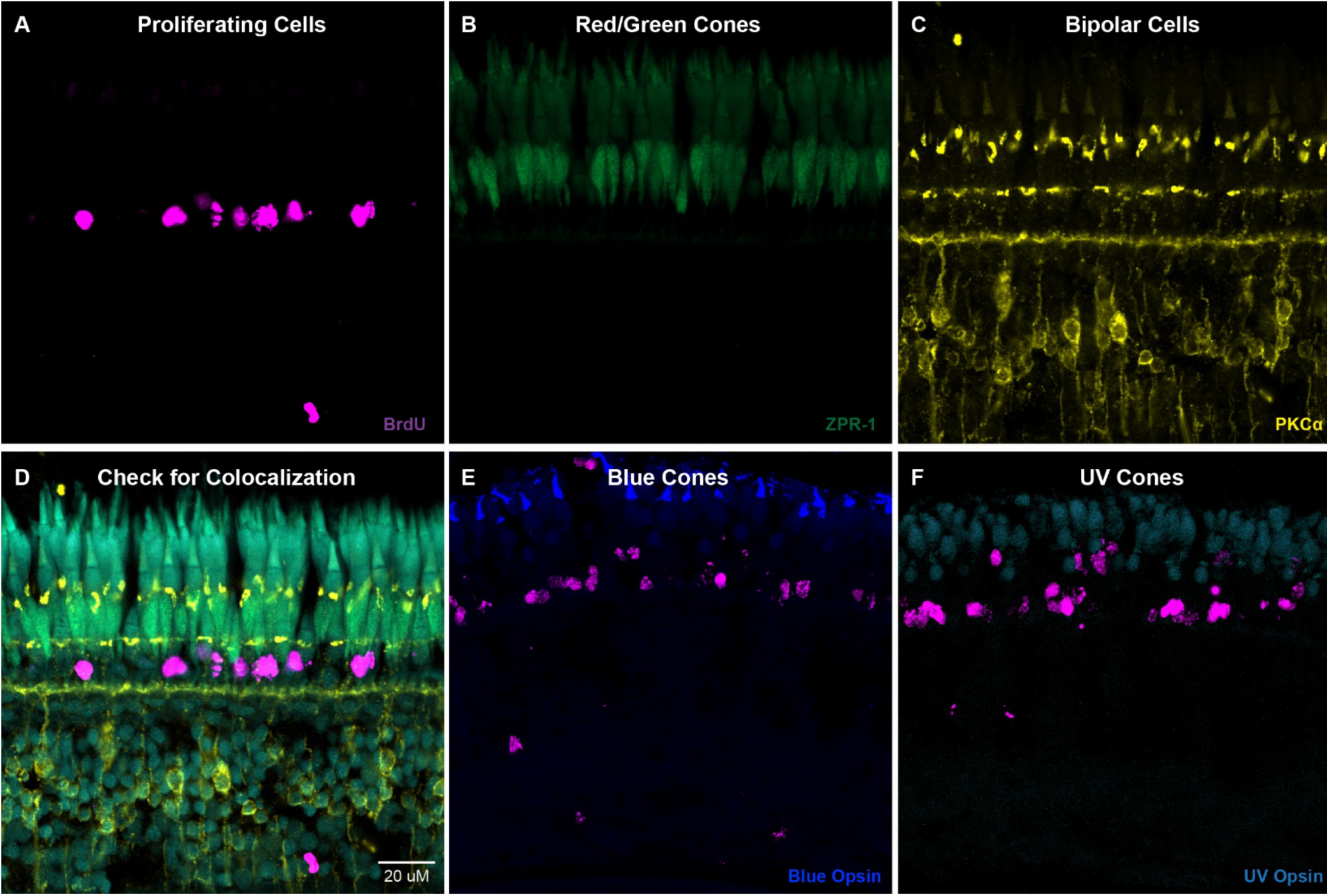
Check for progenitor cell differentiation into other cell types after *prdm1a* knockdown. A) BrdU labelling of P23H retina collected 4 days after *prdm1a* knockdown. B) Immunostaining for red/green cones in P23H retina collected 4 days after *prdm1a* knockdown. C) Immunostaining for bipolar cells in P23H retina collected 4 days after *prdm1a* knockdown. D) Colocalization of A, B, and C show that BrdU-positive cells do not express cone arrestin or *pkcα.* E) BrdU-positive cells do not express Blue Opsin. F) BrdU positive cells do not express UV opsin.

## Discussion

Retinitis Pigmentosa is a chronic retinal degenerative disease characterized by initial degeneration of rod photoreceptors followed by complete degeneration of all photoreceptors through a bystander effect (1, 3, 35–38). Early stages of RP are characterized by night blindness and loss of peripheral vision as rod photoreceptors gradually degenerate (1, 38). Later stages of RP are characterized by complete blindness as cone photoreceptors degenerate via the bystander effect (37, 39). While gene therapy for one variant of RP has been effective, the large number of genes that can cause RP has left many patients with no effective treatment options (4–6). As such, regenerative medicine offers significant hope in preserving and restoring vision to patients with RP (7, 8).

Several studies have investigated regenerative approaches to replace rod photoreceptors and prevent further degeneration of cone photoreceptors as well as replace all photoreceptors to restore vision (8, 40, 41). While not found to occur naturally in mammalian models, Müller glial cells (MGCs) do have the potential to de-differentiate into multipotent progenitor cells, proliferate, and re-differentiate to replace lost retinal cells (42, 43). Regeneration is achieved through MGC activation via *lin28/ascl1* overexpression (44–46). After activation, MGCs de-differentiate into multi-potent progenitor cells, proliferate, and re-differentiate into different retinal cell types (47–49). Several studies have been successful in driving these progenitor cells to re-differentiate into specific populations of cone photoreceptors, bipolar cells (BPCs), amacrine cells (ACs), and retinal ganglion cells (RGCs) in models with retinal injury (45, 47, 50–57). However, rod photoreceptor-specific regeneration has been much more difficult to attain as rod photoreceptors formed in limited numbers, did not successfully integrate into the retina, and/or have only been achieved in a model with no retinal injury (8, 58).

Unlike mammalian models, Zf have the ability to naturally regenerate neurons when injury or disease is detected (9, 12, 14). In acute damage models, MGCs activate, de-differentiate into multipotent progenitor cells, proliferate, re-differentiate into retinal cells, and integrate into the retina (10, 14–18). Many acute damage paradigms ablate groups of retinal neurons (e.g. photoreceptors, outer retinal cells, inner retinal cells) and have identified that progenitor cells derived from MGCs follow fate-specific programs similar to retina formation during development (9, 15, 16, 40, 41, 50, 59–61). While regeneration in acute damaged Zf models stems from MGC activation parallel to mammalian models, regeneration of rod photoreceptors in a Zf model with chronic photoreceptor degeneration stems from retinal progenitor cells found in the ONL (19, 20, 25, 26, 50, 61–65). The mechanisms that drive these progenitor cells to proliferate, differentiate into rod photoreceptors, and develop into mature rod photoreceptors remain unknown. In this study, we investigated these mechanisms and have identified master regulators that control each step of regeneration.

In our P23H Zf model with chronic rod photoreceptor degeneration and regeneration, we have found progenitor cells actively proliferating predominantly throughout the ONL of the retina (Figure 1C). SC-RNA Seq analysis identified 2 progenitor cell populations in our P23H Zf model (Figure 1D). Through HiPlex in-situ hybridization, we were able to identify the RPC cluster as the predominant cluster of progenitor cells found in the ONL (Figure 2D,E). When performing a pulse-chase experiment, we were able to show that these progenitor cells differentiated into rod photoreceptors and degenerated within a span of 1 week (Figure 3D-G). Our Zf model shows continuous degeneration and regeneration of only rod photoreceptors throughout the ventral, dorsal, and central ONL regions of the retina in adult Zf (26). This unique model allows us to study what regeneration would look like in mammalian models with early phases of RP, when cone photoreceptors have yet to begun degenerating and retina remodeling is limited (27, 66, 67). The findings in this study could potentially lead to a higher success rate of permanent rod photoreceptor integration in mammalian models at an early stage of RP.

Monocle3 analysis of the SC-RNA seq data had predicted a trajectory of NPCs differentiating into rod photoreceptors, bipolar cells, or amacrine cells for both WT and P23H datasets (Figure 4A, B). HiPlex in-situ hybridization has shown that this cluster appears to be located in the INL throughout the retina (Figure 2D), while PCNA immunohistochemistry and BrdU incorporation showed very little active proliferation in the INL (Figures 1C, 3C, F). Previous studies have shown that MGCs undergo multiple states between de-differentiation into multipotent progenitor cells and becoming proliferative progenitor cells (50, 68, 69). We believe this cluster could represent non-proliferative multipotent progenitor cells and be a source of replenishment for RPCs in the ONL.

Unique to the P23H dataset were 2 other trajectories that predicted progenitor cell differentiation into rod photoreceptors and maturation of these newly formed rods (Figure 4C, D). In these trajectories, regeneration was divided into 3 main phases: proliferation of progenitor cells, differentiation of progenitor cells into rod photoreceptors, and maturation of newly formed rod photoreceptors. DrivAER analysis predicted a set of master transcription factor regulators responsible for each phase of regeneration. *E2f1, e2f2,* and *e2f3* were important master regulators of progenitor cell proliferation (Figure 5A). E2f1, E2f2, and E2f3 are known to be cell cycle activators that play an important role in progressing proliferation (70–72). Knockdown of these transcription factors in our P23H model resulted in a significant decrease in proliferation (Figure 6C, D). In acute damaged models, *myc/lin28/ascl1* have been found to drive proliferation of NPCs (43, 52). In our SC-RNA seq analysis, these genes were not highly upregulated or unique to progenitor cell populations, suggesting that proliferation in a chronic model undergoes an alternative mechanism than acutely damaged models.

Mass spec analysis of whole retina tissues collected after *e2fs1-3* knockdown revealed a significant downregulation of the targeted proteins as well as proteins involved in retina development and photoreceptor cell development (Figure 7A; Table 3). Furthermore, regulation of transcription through ncRNA processing, rRNA processing, nucleobase transport, and protein trafficking were significantly downregulated while processes involved in RNA and histone methyltransferase were upregulated when compared to the control (Figure 7A; Table 3, 4). Similarly, mitochondrial processes involved with cell division and proliferation were downregulated, while processes involved in negatively regulating autophagy were upregulated (Figure 7A; Table 3, 4). Through mass spec analysis, knockdown of *e2fs1-3* appears to show a decrease in proliferation throughout the retina. As proliferation, and by extension regeneration, is decreased, the increase in different metabolic processes activated indicates that the remaining cells may be undergoing additional stress and are compensating metabolically. Previously, we have shown that with continuous degeneration and regeneration, the retinal environment has a significant increase in oxidative metabolism, glycolysis, and synaptic remodeling when compared to WT (27). We believe that the knockdown of regeneration further elevates the stress of retinal cells throughout the retina, enhancing these changes. DrivAER analysis also predicted *prdm1a* (previously called *blimp1a*) to be the top master regulator driving progenitor cell differentiation into rod photoreceptors (Figure 5). Many studies have shown that rod photoreceptor differentiation during development can be driven with expression of *otx2*, *crx*, and *nrl* (73–75). During development, late progenitor cells can differentiate into either bipolar cells or rod photoreceptors (69). While both cell types are *otx2* positive*, prdm1a* expression has been found to stabilize the progenitor cell fate towards photoreceptors and is found to be upstream of *crx* and *nrl* activity (76, 77). Knockdown of *prdm1a* in our Zf model resulted in a significant reduction of the number of rod photoreceptors as early as 48 hours post knockdown, without depleting the number of BrdU positive cells (Figures 8B, C, 9B, C). At 4 days post knockdown, there was a significant reduction in the number of BrdU-positive cells that had differentiated into rod photoreceptors (Figure 9B’’, D), but these cells did not differentiate into other types of neurons that we could detect (Figure 10). Thus, *prdm1a* knockdown halted RPC differentiation into rod photoreceptors, but did not change the cell fate. This finding is consistent with other studies that claim progenitor cells in the ONL are unipotent for rod photoreceptors (20, 25).

Mass spec analysis of retinas collected after *prdm1a* knockdown revealed a significant downregulation of processes involved in visual perception (Figure 8E Tables 7, 8). Furthermore, genes directly targeted by *prdm1a* as well as processes involved in cell differentiation, neuronal morphogenesis, axon guidance, and embryonic pattern specification were all downregulated (Figure 8E; Tables 7, 8). Altogether, knockdown of *prdm1a* confirms it to be an important transcription factor in rod photoreceptor formation and development. Surprisingly, when *prdm1a* was knocked down, the whole retina saw a downregulation of the negative regulation of the innate immune response (i.e. an upregulation of the innate immune response; Figure 8E; Table 7). As microglia are the only resident innate immune cells found in the retina, it appears that *prdm1a* may play a role in communicating with them. Previous studies have shown that a pro-inflammatory response is necessary for MGC derived regeneration (64, 78, 79). As our model shows regeneration is predominantly from progenitor cells in the ONL, with very little activity from MGCs, prolonged knockdown of *prdm1a* could result in the pro-inflammatory response necessary to activate MGCs.

There was a significantly higher number of mislocalized rhodopsin-expressing cells in the INL in the *prdm1a* knockdowns at 4 days post knockdown as compared to the control (Figure 9B, C, D). These cells may have differentiated from MGC derived NPCs found in the INL (Figure 1C, 8B). However, they may result from a phenomenon paralleling nuclear migration found during retinal development (80, 81). Through this process, progenitor cells move in an apical/basal motion in step with the cell cycle (80, 81). After exiting the cell cycle differentiated neurons have then been known to translocate and migrate towards their typical location (80, 81). Future studies will have to further investigate this possibility.

In this study, we have identified a class of progenitor cells that appear to be responsible for rod photoreceptor regeneration. We have used SC-RNA Seq analysis to predict master regulators responsible for each step of rod photoreceptor regeneration: proliferation of progenitor cells, differentiation of progenitor cells into rod photoreceptors, and maturation of rod photoreceptors. We have validated the roles of the master regulators responsible for the proliferation of progenitor cells and differentiation of progenitor cells into rod photoreceptors with transient gene knockdown studies. While this study identifies how rod photoreceptors are regenerated in a Zf model with chronic retinal degeneration parallel to early stages of RP in mammalian models, there is still a need for the investigation of how rod photoreceptors are able to mature and successfully integrate into the retinal circuitry.

## Materials and Methods

### Animal Husbandry

Zebrafish (Danio rerio) used for this study were all raised, spawned, and maintained on a 14 h light/10 h dark cycle in group housing following standard procedures in the zebrafish community (82). P23H rhodopsin transgenic zebrafish (ZFIN name uth4Tg; RRID:ZFIN_ZDB-GENO-220323-6) were developed in our lab and previously characterized (26–28). Wild-type (WT) AB zebrafish (RRID:ZFIN_ZDB-GENO-960809-7) were purchased from the Zebrafish International Resources Center (ZIRC; Eugene, OR, USA). All Zf used in this study were between 6 months and 1 year of age. Upon random selection for experiments, sex and age of the Zf were noted. While we see no difference between sexes of fish, it should be noted that about 40% of the fish used were males and 60% of the fish used were females. All procedures that required the use of Zf comply with the U.S. Public Health Service policy on humane care and use of laboratory animals and the NRC guide for the Care and Use of Laboratory Animals and have been reviewed and approved by the Institutional Animal Care and Use Committee at the University of Houston under PROTO202100037.

### Single-Cell RNA Sequencing

Single-cell RNA Sequencing and analyses of WT AB and P23H adult retinas were previously characterized and published (27). Datasets are available in GEO (accession number GSE234435) and data are available to explore online at https://www.opt.uh.edu/research/zebrafish/.

FASTQ files were aligned to the zebrafish reference genome (GCA_000002035.4_GRCz11) through 10X Genomics CellRanger (V3.1.2.0) ‘count’ function. Output files were run through Babraham Bioinformatics’ FASTQC (V0.11.9) to check on the quality of the sequences used for analysis. Sequences had a normal distribution of GC content as well as base call accuracy above 99%. Over 93% of reads mapped successfully to the genome. Computational analysis was performed through Texas Advanced Computing Center (TACC) Lonestar5 computing service. Scripts were written in Notepad ++ (V7.6.2) and uploaded to TACC through PuTTY (V0.72).

### Seurat Analysis

Aligned data were run through the Seurat package (V3.1.1) in R (V3.6.1)(83, 84). The WT AB dataset contained an average of 69,277 reads per cell and a median of 1,198 unique transcripts (UMIs) per cell. WT analysis detected a total 13,966 cells, 25,673 genes, and 658 median genes per cell. The P23H dataset contained an average of 39,050 reads per cell and a median of 1,241 UMIs per cell. P23H analysis detected a total of 15,585 cells, 26,375 genes, and 684 median genes per cell. Similar to the previous studies, 20 principle components were used to construct nearest neighbor graphs in the PCA space (27). Louvain clustering and non-linear dimensionality reduction via uniform manifold approximation and projection (UMAP) were performed to visualize and explore the clusters at a 1.2 resolution. Each cluster was assigned a cell type identity based on their expression of cell type-specific genes used in previous studies (27, 28). Violin Plots, Dots Plots, and UMAP cluster maps were all generated through Seurat’s vignettes (V3.2).

### Monocle Analysis

Data analyses from Seurat were run through the Monocle3 package (V0.2.0) in R for both WT and P23H datasets. Monocle3 uses a machine learning algorithm to identify the changes in gene expression needed to go from 1 cell type (cluster) to another cell type (cluster) (29, 30). Plotted as an expression score analysis, clusters that express similar genes are given a higher expression score than other clusters. Monocle3 also provides a pseudotime analysis that allows the user to identify the relative likeliness of trajectories. The lower the pseudotime, the more likely the trajectory will occur relative to other clusters with higher pseudotimes. Predicted trajectories with a high expression score and low pseudotime for rod photoreceptor regeneration were selected for this study.

### DrivAER Analysis

Gene lists for each of the predicted trajectories were run through the DrivAER package (V0.0.2) in Python (V3.9.7). DrivAER is a Python package designed to identify the driving transcriptional programs for a set of genes (31). For the genes involved in the predicted trajectories for rod photoreceptor regeneration, DrivAER analysis provided a list of master regulators believed to be responsible for each network of genes. Through DrivAER analysis, we were able to identify predicted master regulators responsible for different stages of regeneration: proliferation of progenitor cells, differentiation of progenitor cells into rod photoreceptors, and maturation of rod photoreceptors.

### BrdU Intraperitoneal Injections

5-bromo-2-deoxyuridine (BrdU) was freshly prepared in sterile phosphate-buffered saline (PBS; 0.01 M phosphate buffer, 0.138 M NaCl, 0.0027 M KCl, pH 7.4) at a concentration of 5 mg/mL. Each fish was anesthetized in 0.02% Tricaine/MS222 until it was no longer responsive to being picked up with a plastic spoon. The zebrafish was then placed on a mold that exposed its belly while keeping the rest of the body submerged in fish water. It was then intraperitoneally injected with 5 µL BrdU per 0.1 g zebrafish body weight. After injections, the zebrafish was replaced in an isolated tank and kept under observation until further injections or collection.

### Vivo-Morpholinos

Vivo-Morpholinos have previously been shown to effectively target and knock down genes of interest in the retinas of adult zebrafish (28). Translation blocking Vivo-morpholino oligonucleotides were designed and synthesized through GeneTools LLC (Philomath, OR, USA). A control Vivo-morpholino was designed to target EGFP (Table 2). Similar to our previous studies, sterile water was used to make a stock concentration of 200 µM for each Vivo-morpholino (28). For *e2f1, e2f2, and e2f3* targeted knockdown injections, a final working concentration of 15 µM for each Vivo-morpholino was prepared and combined in sterile filtered PBS. For *aurkb* and *prdm1a* targeted knockdown injections, a final working concentration of 45 µM for each Vivo-morpholino was prepared in sterile filtered PBS.

### Intravitreal Injections

Intravitreal injections were performed on adult zebrafish eyes in the same manner as our previous study (28). Each fish was anesthetized in 0.02% Tricaine/MS222 until it was no longer responsive to being picked up with a plastic spoon. A towel was then dipped in fish water and placed inside the lid of a Petri dish. The fish was tucked inside the towel so that only its head was exposed. Under a microscope, a sapphire blade (World Precision Instruments, Sarasota, FL, USA; Catalog #504072) was used to make an incision into the eye between the outer edge of the iris and the pupil. A 32 gauge blunt NEUROS Syringe (Stoelting, Wood Dale, IL, USA; Catalog #53493) was inserted at the site of incision and 1.0 µl of vitreous humor was removed from the eye. After expelling vitreous humor from the needle, 1.0 µL of either the Vivo-morpholino solution of interest or control was injected into the site of incision. The fish was then replaced in an isolated tank and kept under observation until further injections or collection.

### Experimental Paradigms

#### Proliferation

Adult P23H Zf were intravitreally injected with a cocktail of the *e2f1, e2f2,* and *e2f3* Vivo-morpholinos, *aurkb* Vivo-morpholino, or a control Vivo-morpholino. At 24 hours post initial injection (hpii), they were given intraperitoneal injections of BrdU. At 48 hpii, Zf were again given intraperitoneal injections of BrdU. Zf retinas were then collected at 53 hpii. Tissue slices were stained and quantified for the number of BrdU positive cells present per 40x image field of retina. Tissue slices across the ventral, central, and dorsal regions of each retina were quantified.

To determine whether *prdm1a* affected the proliferation of progenitor cells, adult P23H Zf were intravitreally injected with either the *prdm1a* Vivo-morpholino or control Vivo-morpholino. At 24 hpii, Zf were given intraperitoneal injections of BrdU. At 48 hpii, Zf were again given intraperitoneal injections of BrdU. Zf retinas were then collected at 53 hpii. Tissue slices from the ventral, central and dorsal regions of the retina were stained and quantified for both BrdU and rhodopsin labeling.

#### Differentiation

In order to test whether differentiation is affected by *prdm1a* inhibition, adult P23H Zf were first given intraperitoneal injections of BrdU. At 24 hpii, they were again given intraperitoneal injections of BrdU as well as intravitreal injections of either the *prdm1a* Vivo-morpholino or control Vivo-morpholino. Because the half-life of newly formed rods is within 1 week, Zf were maintained for an extra 4 days after the final injection to capture progenitor cell differentiation into rods. Zf were then collected 4 days after intravitreal injections. Tissue slices from the ventral, central, and dorsal regions of the retina were stained and quantified for both BrdU and rhodopsin labeling.

### Immunohistochemistry

Adult zebrafish eyes were collected and fixed in 4% paraformaldehyde (PFA) in 0.1M phosphate buffer (PB), pH 7.5, for 0.5 h at RT. Eyes were then washed 3 times at 5 minute intervals in PB at RT before being placed in 30% sucrose in PB overnight at 4°C. The next day, eyes were embedded in Tissue-Tek O.C.T. compound (Sakura Finetek, Torrance, CA, USA) and stored at -80°C. Retinal sections were cut at a 10 µm thickness and slides were stored at -20°C. Slides from the dorsal, ventral, and center of the retina were used for immunohistochemistry. Slides were washed 3 times at room temperature (RT) in PBS before being treated with blocking solution containing 0.3% Triton X-100 and 5% of the serum of the species in which the secondary antibody was generated (either Donkey Serum or Goat Serum; Jackson ImmunoResearch; West Grove, PA, USA) in PBS for 1 h at RT. Next, slides were treated with primary antibodies diluted in PBS, 0.1% Triton X-100, and 5% serum overnight at RT. The next day, slides were washed 3 times at RT in PBS before being treated with fluorescent secondary antibody diluted in PBS, 0.1% Triton X-100, and 5% serum for 1 h at RT. For nuclear counterstaining, retinal sections were mounted in Vectashield with DAPI (H-1000; Vector Laboratories, Burlingame, CA, USA) and coverslipped. The primary and secondary antibodies used in this study are listed in Table 9. Images were taken using a Zeiss LSM 800 confocal microscope with a 40x objective lens (Thornwood, NY, USA).

**Table 9.** Primary and secondary antibodies used in this study.

| Antibody | Host | Antigen | Source | Catalog Number | RRID | Dilution |
| --- | --- | --- | --- | --- | --- | --- |
| BU-1 | Ms | 5-bromo-2-deoxyuridine (BrdU) | Invitrogen Carlsbad, CA, USA | MA3-071 | AB_10986341 | 1:100 |
| Flag-DDK | Ms | DYKDDDDK | Origene Rockville, MD, USA | TA50011 | AB_2622345 | 1:250 |
| PCNA | Ms | Protein A-Proliferating Cell Nuclear Antigen fusion protein | Abcam Cambridge, MA, USA | ab29 | AB_303394 | 1:100 |
| PCNA | Rb | Synthetic peptide corresponding to Human PCNA aa 200 to the C-terminus | Abcam Cambridge, MA, USA | Ab18197 | AB_444313 | 1:100 |
| PKCα | Rb | Protein Kinase Cα | Millipore, Sigma | P4334 | AB_477345 | 1:400 |
| Retp1 | Ms | Rat Rhodopsin | Novus Biologicals Centennial, CO, USA | NB120-3267 | AB_791922 | 1:200 |
| Zpr1/Fret43 | Ms | Red/Green cone arrestin | ZIRC Eugene, OR, USA | ab174435 | AB_10013803 | 1:10 |
| Cy3 | Gt | Goat Anti-Mouse IgG Fcγ subclass 2a specific | Jackson ImmunoResearch West Grove, PA, USA | 115-165-206 | AB_2338695 | 1:500 |
| Alexa Fluor 488 | Dk | Donkey Anti-Mouse IgG (H+L) | Jackson ImmunoResearch West Grove, PA, USA | 711-545-150 | AB_2340846 | 1:500 |
| Alexa Fluor 488 | Dk | Donkey Anti-Rabbit IgG | Jackson ImmunoResearch West Grove, PA, USA | 711-545-152 | AB_2313584 | 1:500 |
| Alexa Fluor 488 | Gt | Goat Anti-Mouse IgG Fcγ subclass 1 specific | Jackson ImmunoResearch West Grove, PA, USA | 115-545-205 | AB_2338854 | 1:500 |
| Alexa Fluor 647 | Dk | Donkey Anti-Mouse IgG (H+L) | Jackson ImmunoResearch West Grove, PA, USA | 715-605-151 | AB_2340863 | 1:500 |
| Alexa Fluor 647 | Dk | Donkey Anti-Rabbit IgG | Jackson ImmunoResearch West Grove, PA, USA | 711-605-152 | AB_2492288 | 1:500 |
| DAPI |  | Nuclear Counterstaining | Vector Laboratories Burlingame, CA, USA | H-1900-10 |  |  |

### Terminal Deoxynucleotidyl Transferase dUTP Nick End (TUNEL) Labeling

Slides from the dorsal, ventral, and center of the retina were used for TUNEL labeling. Slides were washed 3 times at RT in PBS before being treated with blocking solution containing 0.3% Triton X-100 and 5% Donkey Serum (Jackson ImmunoResearh, West Grove, PA, USA) in PBS for 1 h at RT. Next, slides were treated with the TUNEL reaction mixture containing a 1:9 ratio of Enzyme Solution to Label Solution (provided with In Situ Cell Death Detection Kit 12156792910; Millipore Sigma, Burlington, MA, USA). After 1 h of incubation at RT in the dark, slides were washed 3 times at RT in PBS. For nuclear counterstaining, retinal sections were mounted in Vectashield with DAPI and coverslipped. Slides were immediately imaged using a Zeiss LSM 800 laser scanning confocal microscope.

### Multiplex Fluorescent In-Situ Hybridization

Adult zebrafish eyes were collected and fixed in 4% paraformaldehyde (PFA) in 0.1M phosphate buffer (PB), pH 7.5, for 24 h at RT. Eyes were then washed 4 times at 15 minute intervals in PB at RT before being placed in 30% sucrose in PB overnight at 4°C. The next day, eyes were embedded in Tissue-Tek O.C.T. compound and stored at -80°C. Retinal sections were cut at a 20 µm thickness and slides were stored at -80°C. Through RNAscope HiPlex (Advanced Cell Diagnostics, Newark, CA, USA), we generated in-situ probes for specific genes of interest (Table 10). Sections were dehydrated in 50% ethanol (ETOH), 70% ETOH, and 100% ETOH for 5 minutes at each concentration at RT. Tissues were then treated with protease (provided with HiPlex kit 324419) for 30 minutes at RT followed by a 5 minute wash in PBS at RT. They were then treated with the 1X Target Retrieval Reagent (provided with HiPlex kit 324419) for 5 minutes at 95°C. Following manufacturer’s instructions, probes were applied to each tissue sample and allowed to hybridize for 2 hours before going through 3 amplification treatment steps that each last for 30 minutes (85). After final amplification, tissues were treated with detection solution for 15 minutes and counterstained with DAPI before being mounted. Slides were immediately imaged using a Zeiss LSM 800 laser scanning confocal microscope.

**Table 10.**
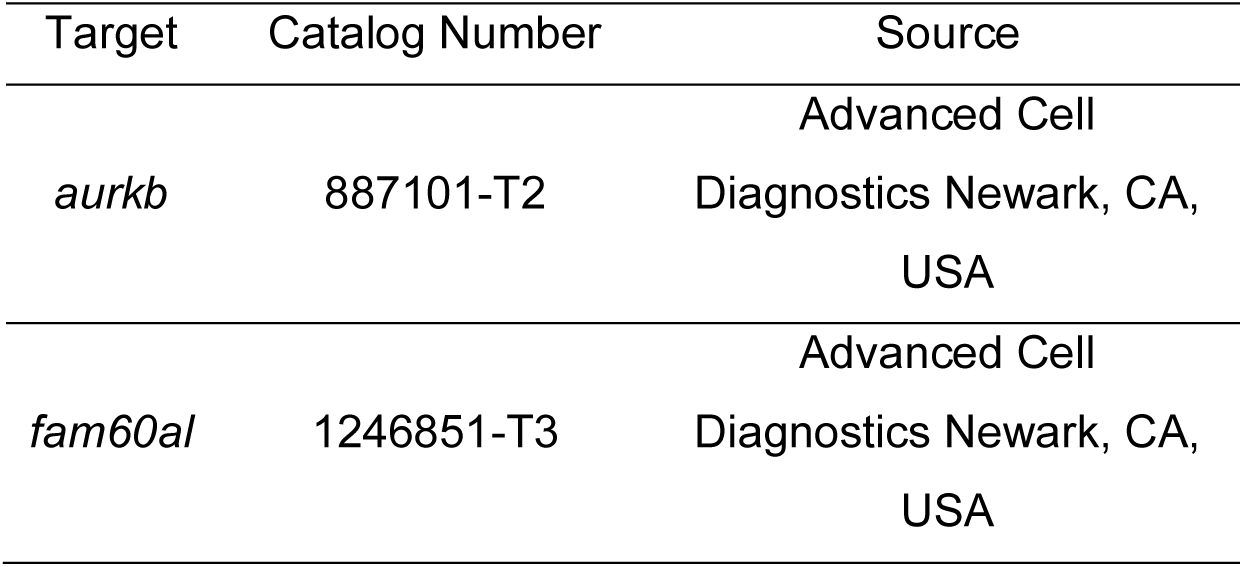
HiPlex probes used in this study.

### Quantification

For each retina used for immunohistochemistry, 9 sections throughout the dorsal, central, and ventral retina were labeled, imaged, and analyzed. For each retina used for TUNEL staining, 6 sections throughout the dorsal, central, and ventral retina were labeled, imaged, and analyzed. An image sequence (6.63 µm total thickness divided into 17 sections, each 0.39 µm thick) was taken with a 40X objective lens at 1x confocal zoom (160µm x 160 µm image field). A maximum intensity projection of each sequence was made and imported into FIJI (ImageJ 1.53t). The image was then split by each channel (i.e., red, green, blue). For each channel, a threshold was adjusted to capture only the signal. After applying the threshold, a binary image mask showed the signal as white and background as black. When measuring expression levels, for each image the ratio of white pixels (signal) to total pixels was calculated. To assess differentiation of progenitor cells into rods, the ‘colocalization’ plugin in FIJI was used to assess how many cells expressed both BrdU and rhodopsin (RetP1 immunolabeling). To measure TUNEL-positive cells, the number of regions of interest identified by the threshold was counted. For each experimental animal (biological replicates), the mean data from the 9 imaged sections were compared for statistical analyses. In Graphpad Prism 9, a two-tailed t-test was performed to compare signals between the control and injected fish. For the BrdU-pulse chase and proliferation knockdown experiments, an ANOVA analysis followed by Tukey’s post-hoc test was performed to compare quantifications between different conditions. A p-value <0.05 was considered significantly different.

### Liquid Chromatography Mass Spectrometry Proteomics

After carrying out the experimental paradigms described earlier, retinas were collected, flash frozen in liquid nitrogen, and stored at -80°C until prepared for mass spectrometry. For each condition, two retinas from separate animals were processed separately and treated as biological replicates. The proteomic sample preparation was performed by following the Sample Preparation by Easy Extraction and Digestion (SPEED) procedure previously described by Doellinger, et al. (86). Briefly, 10 µl of trifluoroacetic acid were added to each retina tissue sample and vortexed until all samples were dissolved. Next, 100 µl of 2M tris base was added, followed by a mixture of TCEP (10 mM) and CAA (40 mM). The samples were heated at 95°C for 5 minutes before adding 500 µl of HPLC grade water. 3 µg of proteins from each sample were transferred to a 96-well plate with 30 µg of SeraSil-Mag magnetic beads (28967388, Cytiva, Marlborough, MA, USA) per well. An SP3 protocol and on-bead digestion using Trypsin (w/w 1:50) were used to generate peptide samples (87). 100 ng of peptides were loaded onto Evotip and analyzed using an EVOSEP One LC (EVOSEP, Odense, Denmark) connected to a timsTOF Pro2 (Bruker Daltonics, Bremen, Germany). The standard 60 sample per day method (21-minute gradient) was used with a C18 column (EV1109, EVOSEP). A data-independent acquisition parallel accumulation-serial fragmentation (diaPASEF) scheme with 16 m/z and ion mobility windows was used for mass spec (MS) data acquisition. The electrospray voltage was 1.5 kV, and the ion transfer tube temperature was 180°C. Full MS scans were acquired over the range m/z of 100-1700. The collision energy was ramped linearly as a function of the mobility from 20 eV at 1/K0 = 0.6 V•s/cm^2^ to 59 eV at 1/K0 = 1.6 V•s/cm^2^.

### MS Data Analysis

The software DIANN version 1.8 was used with default settings for peptide and protein identification and quantification from diaPASEF data (88). Data were searched against the UniProt-SwissProt *Danio rerio* database (Taxon ID 7955 downloaded on 8/27/2024; 3338 entries). Cysteine carbamidomethylation was listed as a fixed modification and methionine oxidation and acetylation as variable modifications. The false discovery rate (FDR) was controlled at <1% at both peptide and protein levels. For each experimental condition performed, duplicates were run through MS analysis (Supplementary Table 2). Expression levels were averaged. A ratio of the experimental condition to its proper control was used to identify proteins that were greater than 3 log2 folds or less than 0.5 log2 folds. Proteins within each threshold were run through the Cytoscape (V3.10.2) plugin ClueGo (V2.5.10) to identify the GO Biological Processes upregulated or downregulated for each dataset.

## Acknowledgements

This work was supported by a grant from the William Stamps Farish Fund, RF1MH120016, core grants P30EY028102 and P30EY007551, and the University of Houston College of Optometry. ES was supported by F31EY034793. JW is supported by NIH grants R01NS088353 and R21NS113068 and the Amy and Edward Knight Fund of the UTHSC Senator Lloyd Bentsen Stroke Center. GQ is supported by the National Science Foundation grant DMR2005199.

The authors would like to thank Dr. Steven W. Wang for guidance with intraperitoneal injections and Dr. Stephan Tetenborg for helpful discussions. The authors would also like to thank Dr. David Hyde for providing cone opsin antibodies. The authors would also like to thank the Division of Research, University of Houston, for purchase of the Evosep One system; Mr. Stan Stearns for his generous gift and Valco Instruments Company Incorporated (VICI) for the donation of the Agilent Bravo AssayMap System.

## Author contributions

Conceptualization: ES, JO; Methodology: ES, AS, GQ; Validation: ES, AS; Formal analysis: ES, AS, AA; Investigation: ES, ST, AS, AA, HW, JW, GQ; Resources: ES, AS, JO; Data curation: ES, AS; Writing-original draft: ES; Writing-review and editing: ES, AS, JO, HW, JW, AA, ST, GQ, CC; Visualization: ES, AS, AA, JO; Supervision: JO, JW, CC; Project administration: JO; Funding acquisition: JO, ES, JW, CC, GQ.

## Data and materials availability

The P23H rhodopsin transgenic zebrafish strain Tg (rho:MmuRho_P23H-FLAG)^uth4^ is available for research purposes from the authors with a Material Transfer Agreement with the University of Texas Health Science Center at Houston. Data from this study can be explored at https://www.opt.uh.edu/research/zebrafish/. Raw data used in this study have been deposited in GEO under Project Numbers GSE234435 and GSE234661.

---

**Supplementary Figure 1.**
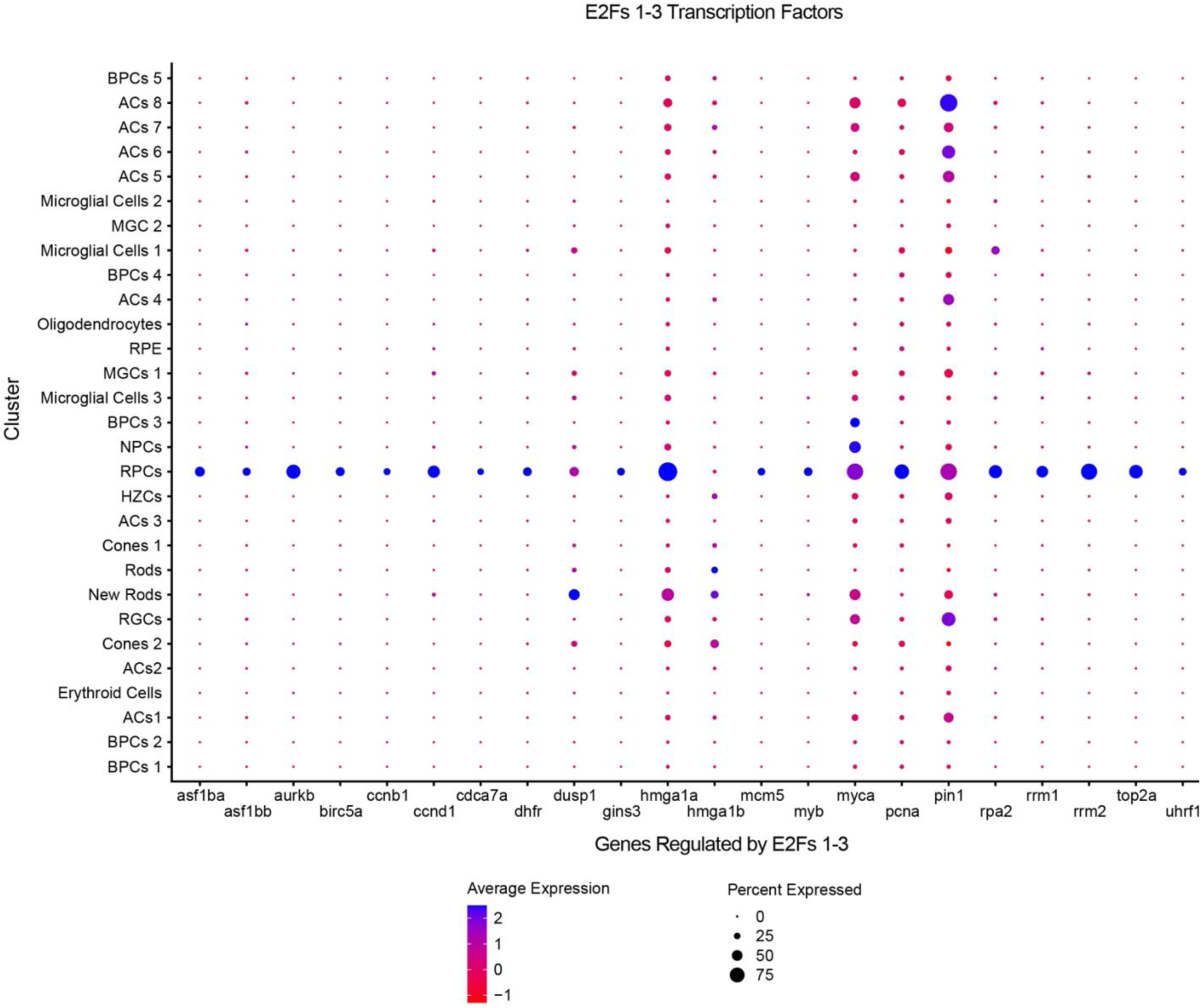
Dotplot analysis of the trajectory genes controlled by *e2f1, e2f2,* and *e2f3*. The list of genes from the trajectory that were predicted to be controlled by the *e2f’s 1-3* has been illustrated in a Dotplot to show their average expression levels (color scale) and fraction of cells expressing the gene (dot size) within each cluster of the P23H dataset.

**Supplementary Figure 2.**
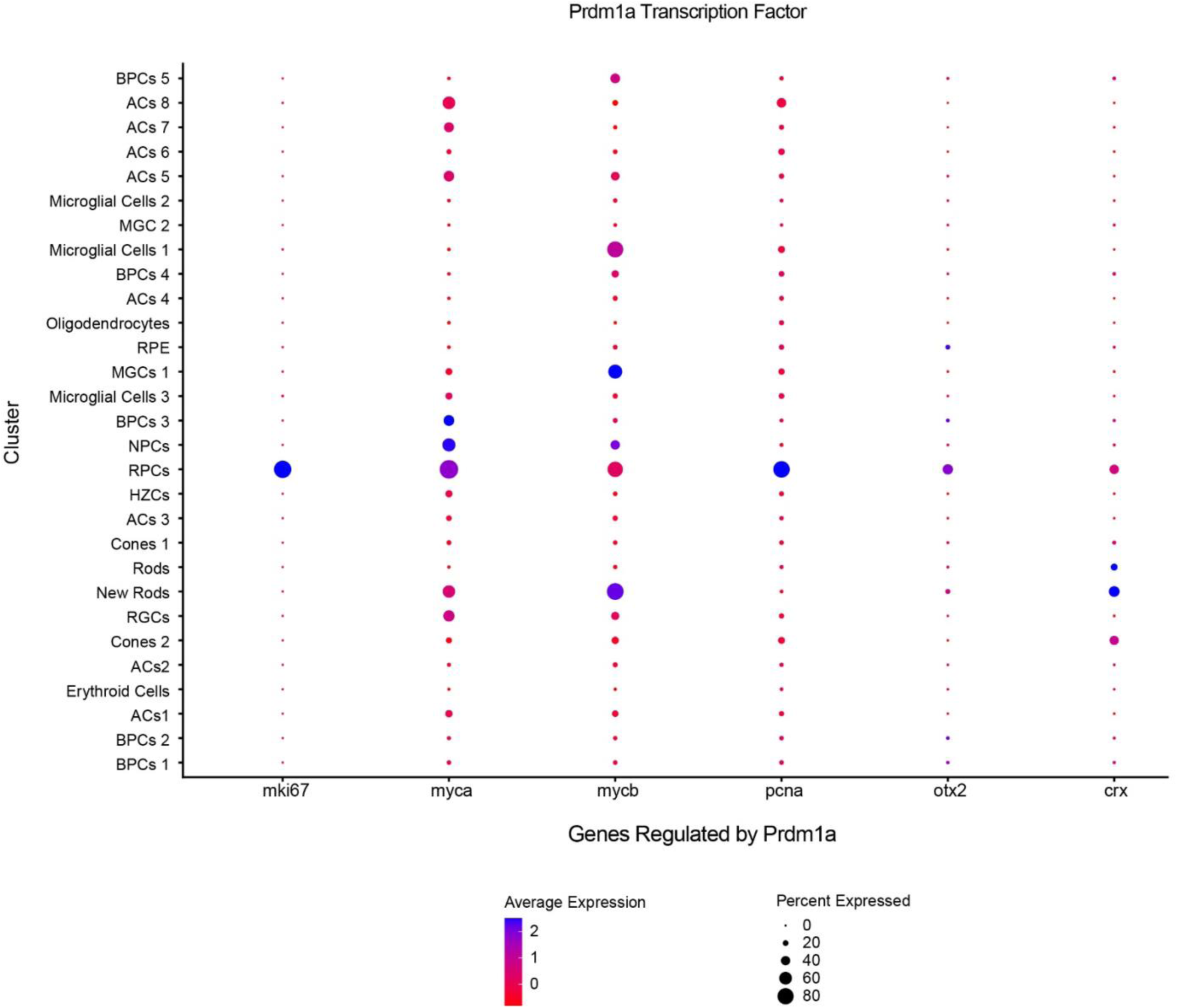
Dotplot analysis of the trajectory genes controlled by *prdm1a*. The list of genes from the trajectory that were predicted to be controlled by *prdm1a* has been illustrated in a Dotplot to show their average expression levels (color scale) and fraction of cells expressing the gene (dot size) within each cluster of the P23H dataset.

**Supplementary Figure 3.**
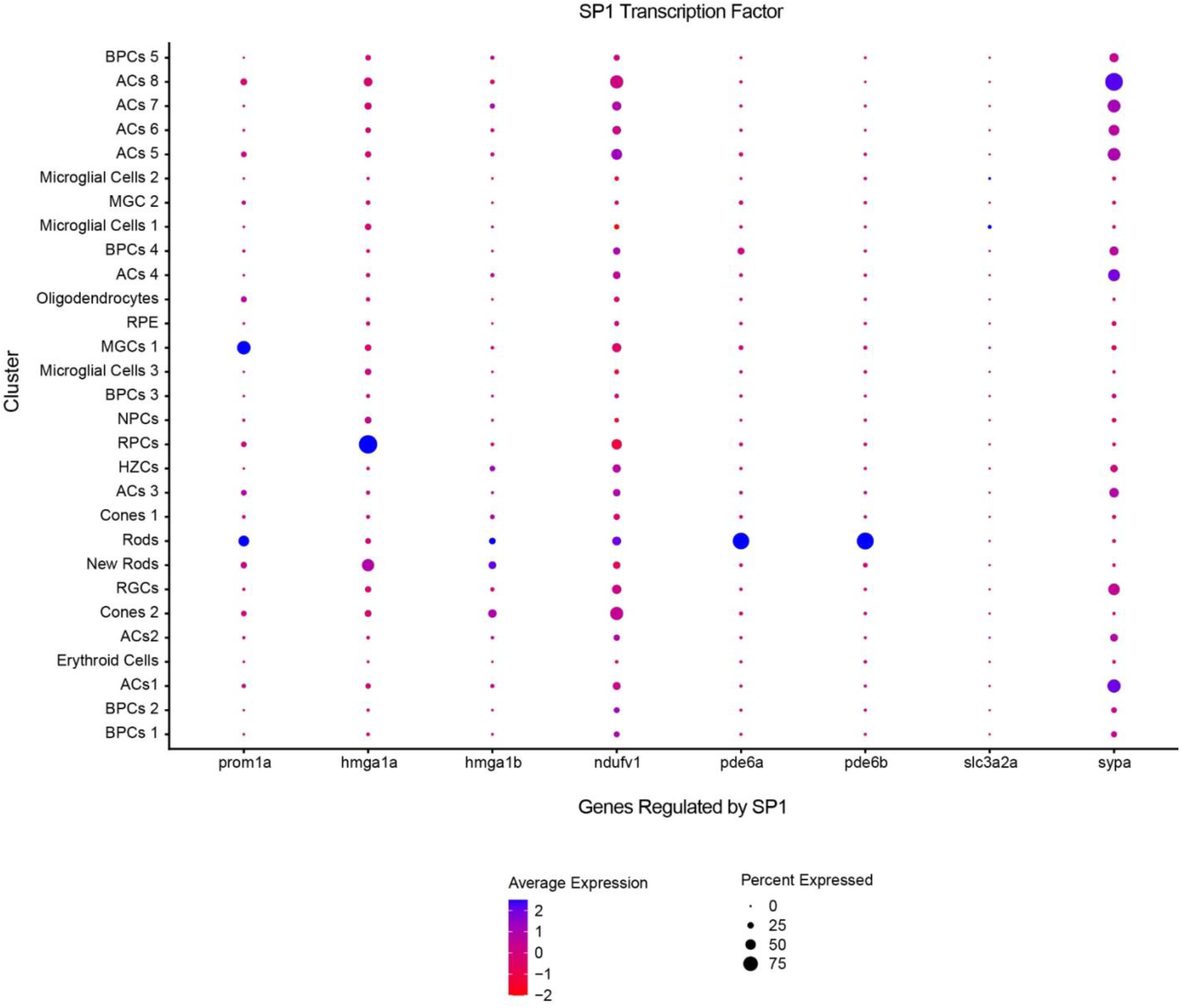
Dotplot analysis of the trajectory genes controlled by *sp1*. The list of genes from the trajectory that were predicted to be controlled by *sp1* has been illustrated in a Dotplot to show their average expression levels (color scale) and fraction of cells expressing the gene (dot size) within each cluster of the P23H dataset.

